# Cellular dynamics across aged human brains uncover a multicellular cascade leading to Alzheimer’s disease

**DOI:** 10.1101/2023.03.07.531493

**Authors:** Gilad Sahar Green, Masashi Fujita, Hyun-Sik Yang, Mariko Taga, Cristin McCabe, Anael Cain, Charles C. White, Anna K. Schmidtner, Lu Zeng, Yangling Wang, Aviv Regev, Vilas Menon, David A. Bennett, Naomi Habib, Philip L. De Jager

**Affiliations:** Edmond & Lily Safra Center for Brain Sciences, The Hebrew University of Jerusalem, Jerusalem, Israel; Center for Translational & Computational Immunology, Department of Neurology and Taub Institute for Research on Alzheimer’s Disease and the Aging Brain, Columbia University Medical Center, New York, New York, USA; Harvard Medical School, Boston, MA; Center for Alzheimer Research and Treatment, Department of Neurology, Brigham and Women’s Hospital, Boston, MA; Broad Institute of MIT and Harvard, Cambridge, Massachusetts, USA; Klarman Cell Observatory, Broad Institute of MIT and Harvard, Cambridge, Massachusetts, USA; Rush Alzheimer’s Disease Center, Rush University Medical Center, Chicago, Illinois, USA

**Author notes:** Equal contribution (Masashi Fujita, Hyun-Sik Yang). Equal contribution (Vilas Menon, David A. Bennett, Naomi Habib, Philip L. De Jager).

## Abstract

Alzheimer’s Disease (AD) is a progressive neurodegenerative disease seen with advancing age. Recent studies have revealed diverse AD-associated cell states, yet when and how they impact the causal chain leading to AD remains unknown. To reconstruct the dynamics of the brain’s cellular environment along the disease cascade and to distinguish between AD and aging effects, we built a comprehensive cell atlas of the aged prefrontal cortex from 1.64 million single-nucleus RNA-seq profiles. We associated glial, vascular and neuronal subpopulations with AD-related traits for 424 aging individuals, and aligned them along the disease cascade using causal modeling. We identified two distinct lipid-associated microglial subpopulations, one contributed to amyloid-β proteinopathy while the other mediated the effect of amyloid-β in accelerating tau proteinopathy, as well as an astrocyte subpopulation that mediated the effect of tau on cognitive decline. To model the coordinated dynamics of the entire cellular environment we devised the BEYOND methodology which uncovered two distinct trajectories of brain aging that are defined by distinct sequences of changes in cellular communities. Older individuals are engaged in one of two possible trajectories, each associated with progressive changes in specific cellular communities that end with: (1) AD dementia or (2) alternative brain aging. Thus, we provide a cellular foundation for a new perspective of AD pathophysiology that could inform the development of new therapeutic interventions targeting cellular communities, while designing a different clinical management for those individuals on the path to AD or to alternative brain aging.

## Introduction

The molecular characterization of the aging human brain has expanded dramatically over the past decade. Platforms for high-throughput measurements of molecular features have become available and have converged with the development of analytic methods capable of providing new insights into the growing mass of data from bulk tissue samples^1, 2^. These studies have demonstrated that with a statistically well-powered sample size, comprehensive analyses of autopsied brain samples can yield reproducible molecular insights into Alzheimer’s disease (AD)^3^. However, bulk profiling of the brain parenchyma loses much of the information embedded in the intricate cellular architecture of the brain structures affected by AD. Single-cell and single-nucleus RNA-seq^4–16^ have begun to offer a different perspective, underscoring the cellular perturbations that are part of the sequence of events leading to AD^17^. Currently, it appears that^4–15^: (1) changes in expression programs of multiple cell types, including glial, neuronal and vascular cells, are associated with AD, (2) within each cell type only specific subpopulations are likely to be actively involved in AD pathogenesis, and (3) AD-related expression programs changes are coordinated across multicellular communities of various cell subpopulations^4^.

The question before us is therefore clear: can we robustly identify the cell populations and environments involved in AD? Untangle the process of AD from the process of brain aging? And prioritize those with a potentially causal role at each step of the disease cascade? One major challenge in addressing this question is accurately aligning individuals along a timeline of disease progression, especially in the preclinical asymptomatic phase, where AD pathologies are present but there is no cognitive decline^18^. Previous studies have avoided this challenge by using a case/control design, focusing on extremes, such as advanced stages of AD, to find cell populations associated with the disease. Moreover, the heterogeneity of clinical symptoms and pathological manifestation in older individuals, while increasingly appreciated, were not adequately addressed in previous studies. To overcome these challenges, we aimed to profile the diversity of cellular environments and their dynamics along the entire cascade of the disease and along non-AD brain aging, in a large collection of prospectively collected aged brains with detailed ante- and post-mortem phenotypic characterization. Such data will address key open questions regarding the cellular cascade(s) leading to AD, the cellular drivers at each disease stage, and the differences between AD and the process of brain aging.

Specifically, we leverage two community-based prospective cohort studies of cognitive aging, the Religious Order Study and the Rush Memory and Aging Project (ROSMAP)^19, 20^, that capture a wide clinicopathologic spectrum of cognitive aging and AD with extensive ante- and post-mortem characterization. We built a cellular atlas from RNA profiles of 1.64 million nuclei from the dorsolateral prefrontal cortex (DLPFC, BA9) of 465 ROSMAP participants, out of which 424 participants were used in downstream analyses. We devised a new computational framework, BEYOND, to align individuals along cellular cascades and capture the diversity of cellular environments and their dynamics along distinct trajectories (**Fig. 1a**). The ROSMAP dataset, our study’s design and the BEYOND approach enabled us to address the key goal of proposing a sequence of cellular events leading to AD and untangling the processes of AD and aging. Our sample size provides the necessary statistical power to investigate the cellular landscape of the aging human brain while accounting for human heterogeneity and to reveal distinct trajectories of cellular change. Along one of these trajectories we are able to propose a statistically rigorous cellular-molecular pseudo-temporal cascade that leads to AD using cross-sectional data derived from autopsy samples. Finally, our detailed cellular atlas and disease trajectories prioritize distinct microglial and astrocytic populations predicted to drive different stages of the causal chain leading to AD, highlighting important new therapeutic targets.

**Figure 1:**
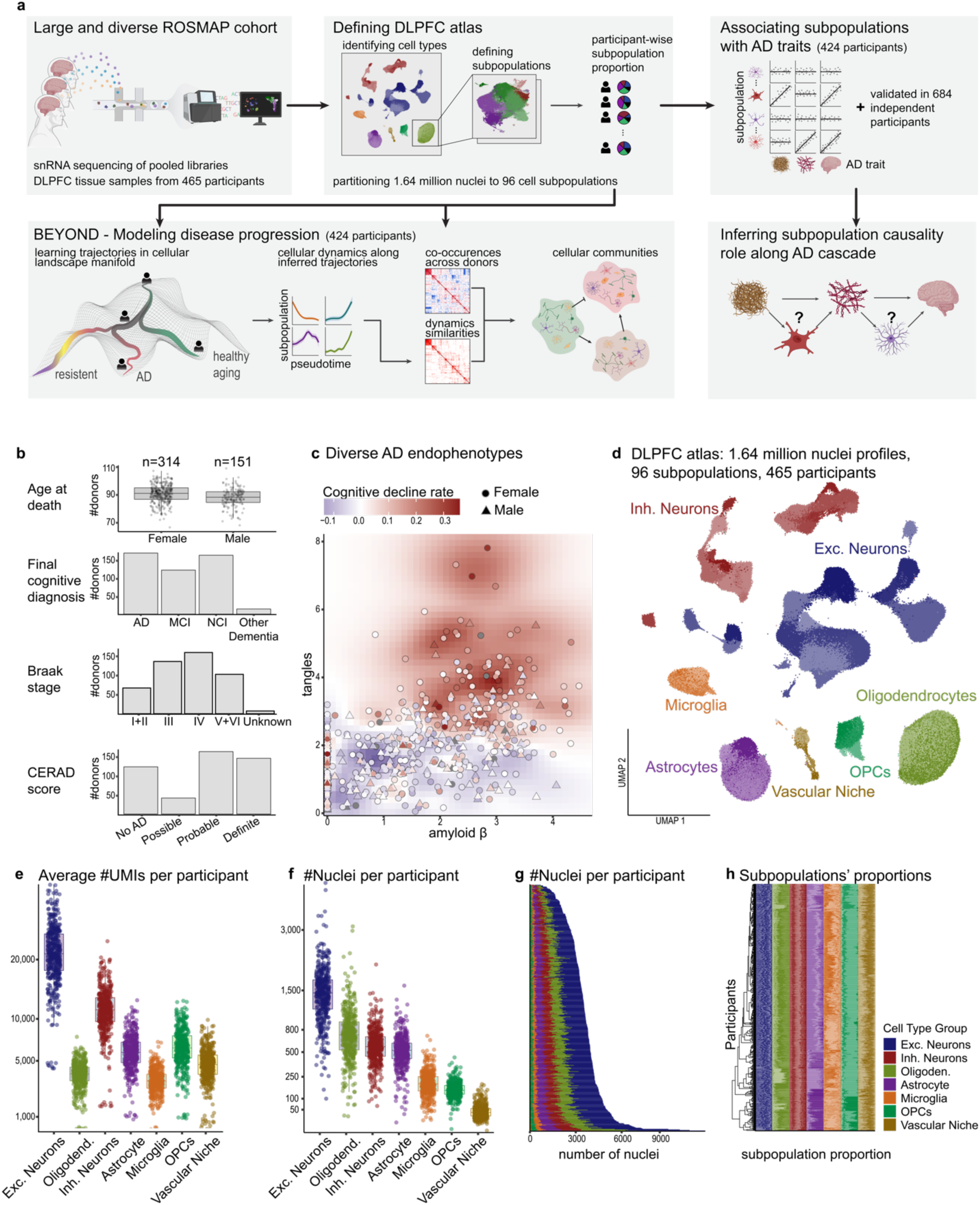
Cellular atlas of the human aged DLPFC in 465 older individuals. (**a**) Overview of experiment and analysis steps. (**b**) Clinicopathologic characteristics of the 465 participants. Additional details are present in **Supplementary Table 1**. (**c**) Distribution of quantitative A*β* and tau measures among the participants. The distribution of the rate of cognitive decline is denoted in color, and the shape denotes the sex of each individual. (**d**) UMAP embedding of 1,638,882 single-nuclei RNA profiles from the DLPFC brain region of 465 participants, color-coded by 96 cell subpopulations. Major cell types are noted. (**e-f**) Quality measures of the atlas showing (e) the average number of UMIs per cell type in each participant and (f) the number of nuclei per cell type in each participant; Dots: individual participants. Box: 25th and 75th percentiles; line: median. (**g**) Number of cells and cell types. Cumulative number of nuclei profiled per participant from each cell type, color-coded by the major cell types. **(h)** Proportions of cell subpopulations. Stacked barplot of cell subpopulation proportions per participant within each major cell type, color-coded by cell type.

## Results

### A high-resolution cell atlas of the aged and AD dorsolateral prefrontal cortex

To generate a cell atlas capturing the cellular heterogeneity in the aged cortex, we performed single nucleus RNA-seq (snRNA-seq) on nuclei isolated from frozen DLPFC tissue samples of 465 ROSMAP participants^19, 20^ (**Fig. 1a**). These two longitudinal studies of cognitive aging enroll non-demented participants and include annual ante-mortem neuropsychological evaluations, prospective brain collection at the time of death, and characterization by a structured neuropathologic assessment. The profiled 465 participants represent a relatively random sample of the deceased ROSMAP participants, and, therefore, they span a wide spectrum of clinicopathologic stages of AD and brain aging (**Fig. 1b-c, Supplementary Fig. 1a, Supplementary Table 1**). The participants average age of death was 89±6.8 years, and 67.5% are females. At the time of death, 62% of the participants fulfill a pathologic diagnosis of AD using the NIA Reagan criteria^21, 22^, while 36% fulfill a clinical AD dementia diagnosis, and 26% show mild cognitive impairment (MCI, **Fig. 1b**). To reduce batch effects, we used a strategy of profiling pools of unsorted nuclei from 8 participants, in two technical replicates, and we assigned cells back to participants by genotyping polymorphic sites in the RNA sequence data and compared it to their reference genome (using *demuxlet*^23^, **Methods, Fig. 1a**). cDNA libraries underwent a detailed quality control and filtering pipeline, including automated cell-type classification, cell-type-specific identification of low-quality nuclei and doublet removal (**Methods, Supplementary Fig. 1**). Ultimately, we retained 1,638,882 high-quality nuclei profiles from 465 participants (**Fig. 1d-h**). We captured all expected brain cell types, and clustered nuclei into 96 known and novel cell subpopulations (**Fig. 1d-h**), each of which is characterized by a distinct expression profile, marker genes and enriched pathways (**Fig. 2, Supplementary Fig. 2-4**). The full annotation of subpopulations, genes and pathways is found in **Supplementary Table 2**.

**Figure 2:**
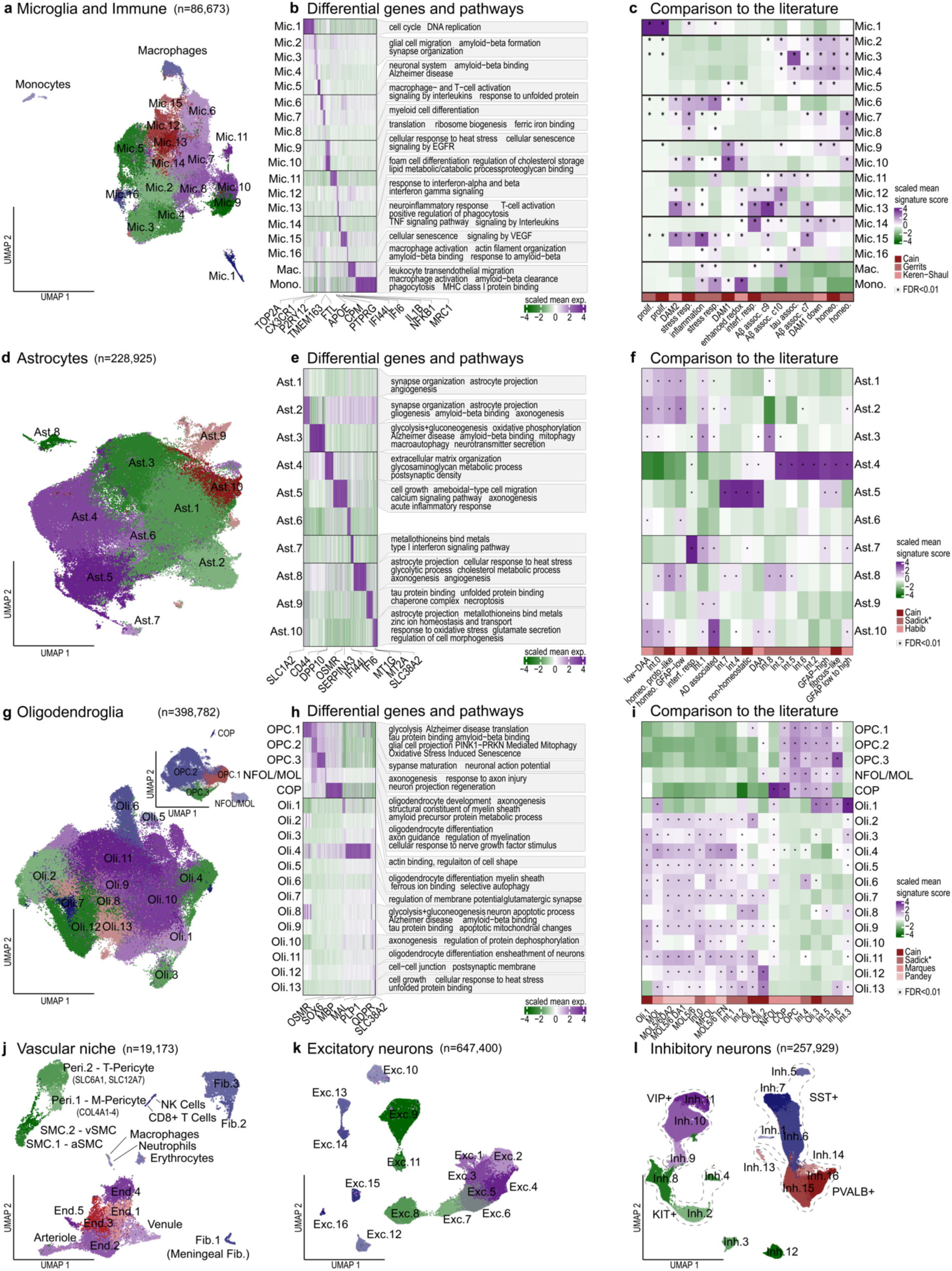
Diversity of cell subpopulations in the aged DLPFC. (**a**) Diversity of microglial subpopulations. UMAP embeddings of single nuclei profiled by snRNA-seq, colored by clusters. (**b**) Distinct gene signatures and associated pathways across microglial subpopulations. Left: differentially expressed genes (columns) for different microglial subpopulations (rows). Color scale: Scaled mean expression per cluster. Right: representative enriched unique pathways per subpopulation. (**c**) Comparison of microglial subpopulations to previous signatures. Scaled mean signature score of published microglial gene signatures (columns) within each microglial subpopulation (rows)^4, 8, 16^. (*) = Significantly enriched signatures (U-test, FDR<0.01). (**d-i**) Diversity of astrocytes and oligodendrocytes. Subpopulations of astrocytes (**d**) and oligodendrocytes (**g**). UMAP embedded nuclei, colored by cluster. Distinct gene signatures and associated pathways of astrocytes (**e**) and oligodendrocytes (**h**), presented as in (b). Comparison of published signatures to subpopulations of astrocytes (**f**) and oligodendrocytes (**i**), presented as in (c). Astrocyte and oligodendrocyte signatures from the Sadick dataset^5^ are an integrated atlas including the Mathys, Grubman and Zhou datasets^6, 10, 14^. (**j**,**k**,**l**) Diversity of vascular and neuronal cells. Subpopulations of (j) vascular niche cells, (k) excitatory neurons and (l) inhibitory neurons. UMAP embedded nuclei, colored by cluster.

At the subpopulation level, microglial nuclei profiles were partitioned into 16 subpopulations (**Fig. 2a-c, Supplementary Fig. 2a-c**, n=86,673 nuclei), including proliferative (Mic.1), surveilling (Mic.2-5; expressing CX3CR1), reacting (Mic.6-8; TMEM163), enhanced-redox (Mic.9-10; FLT1), stress response (Mic.11; NLRP1, TGFBR1, upregulating genes of heat response, cellular senescence and NLRP1 inflammasome), interferon response (Mic.14, IFI6), inflammatory (Mic.15; CCL3/4, NFKB1, NLRP3), SERPINE1 expressing (Mic.16) and lipid-associated (Mic.12-13; APOE) subpopulations. The lipid-associated Mic.12 and Mic.13 both expressed the AD risk genes APOE and GPNMB, with Mic.13 also expressing high levels of SPP1 and TREM2 compared to other subpopulations (**Supplementary Table 2**). These annotations correspond to and further refine previous reports (**Fig. 2c, Supplementary Fig. 3a**). Specifically, Mic.13 and Mic.15 (and to a lower extent also Mic.12) expressed the highest level of mouse-derived DAM2 signature^16^; Mic.12 and Mic.13 expressed the human A*β*-associated microglia signatures^8^; while Mic.15 highly expressed the human inflammation and stress signatures^8^.

Astrocytes were partitioned into 10 subpopulations (**Fig. 2d-f, Supplementary Fig. 2d-f**, n=228,925 nuclei) - homeostatic-like (Ast.1-2), enhanced-mitophagy (Ast.3; PINK1), reactive-like Ast.4 (GFAP, ID3) and Ast.5 (GFAP, SERPINA3, OSMR), interferon-responding (Ast.7; IFI6), and stress response (Ast.8-10): Ast.8, expressing heat stress and DNA damage, calcium, and sterol metabolism genes; Ast.9, expressing heat and oxidative stress response, tau binding and necroptosis genes; and Ast.10 (SLC38A2), expressing oxidative stress and ROS, metallothioneins and zinc ion homeostasis genes. Our astrocyte annotation expands previous reports from healthy and AD human brains, and it links mouse and human signatures^4, 5, 24^: Ast.4 to fibrous-like astrocytes^4^, Ast.5 to mouse disease-associated astrocytes^24^ (DAA), Ast.7 to interferon responding astrocytes^4^, and Ast.10 to human AD elevated astrocytes^4^ (**Supplementary Fig. 3b**).

Oligodendrocyte lineage cells were partitioned into 13 subpopulations of mature oligodendrocytes (Oli.1-13; **Fig. 2g-i, Supplementary Fig. 2g-i, Supplementary Fig. 3c**, n=398,782 nuclei), such as the stress response Oli.13 (SLC38A2), 3 subpopulations of oligodendrocyte precursor cells (OPC.1-3), one committed oligodendrocyte precursor (COP) subpopulation and one newly formed oligodendrocytes (NFOL) subpopulation. The newly discovered diversity of OPCs are of particular interest, and included an enhanced-mitophagy subpopulation (OPC.1; PINK1), which had higher expression of AD risk genes (e.g. APOE, CLU), and an axon projection/regeneration associated subpopulation (OPC.3; SERPINA3, OSMR).

Within the vascular niche (**Fig. 2j, Supplementary Fig. 4a-d**, n=18,496), we identified all expected cell classes, including arterial, venous, and capillary endothelial cells, arterial- and ventricular-smooth muscle cells (SMCs), pericytes, and meningeal and perivascular fibroblasts, which aligns with and expands recent reports^7, 15^ (**Fig. 2j, Supplementary Fig. 4d**). For example, we identified two distinct pericyte subpopulations, Peri.1 and Peri.2, aligning with the recently described extracellular matrix-(M) and transporter (T) pericytes^7^ (respectively). We revealed novel diversity within the capillary endothelial cells (End.1-5), such as End.3 with higher expression of extracellular matrix genes and End.5 expressing heat- and oxidative stress response, tau binding and necroptosis, as well as AD risk genes (e.g. APP and ADAM10).

On the neuronal side, we identified the full range of major classes and subtypes, including 16 subtypes of excitatory neurons (Exc.1-16, n=647,400, **Fig. 2k, Supplementary Fig. 4e-g**), matching previous annotations of neocortical neuronal diversity across the different cortical layers with additional diversity within some layers. We also identified 16 subtypes of inhibitory neurons (Inh.1-16, n=257,929, **Fig. 2l, Supplementary Fig. 4h-j**), matching the major classes expressing distinctive neuropeptides, such as somatostatin (SST, Inh.5-7), parvalbumin (PVALB, Inh.13-16), vasoactive intestinal polypeptide (VIP, Inh.9-11) and KIT (Inh.2, Inh.4 and Inh.8).

### Glial and neuronal subpopulations associated with AD-related traits

To identify subpopulations associated with AD, we focused on three quantitative AD traits available for ROSMAP participants: measures of (1) neocortical amyloid-*β* (A*β*), (2) neocortical tau, and (3) the rate of cognitive decline prior to death measured from up to 20 years of annual neuropsychologic profiles (see **Methods** for the full description of these traits). These outcomes have more statistical power compared to categorical AD diagnosis measures and enabled the assessment of the continuum of AD clinicopathologic severity. For the association and downstream analysis, we only included individuals with a sufficient number of high-quality nuclei that are confidently assigned to them and had whole genome sequence data (**Methods**), reducing the sample size to 424 participants (referred to as the *Discovery sample*, **Supplementary Fig. 1b**).

For our *Discovery analysis*, we tested the association between each of the three AD-related traits and the proportions of all subpopulations (calculated *within* each major compartment by linear regression, while controlling for age, sex, post-mortem interval, and library quality, FDR<0.05, **Fig. 3b, Methods, Supplementary Table 3**). The strongest associations were found for the two lipid-associated microglial subpopulations, Mic.13 and Mic.12 (**Fig. 2a-c**). The strongest signal was for Mic.13 (APOE^+^TREM2^+^, with the highest expression of the DAM2 signature^16^), which was present in higher proportion in individuals with greater neocortical A*β* burden, tau burden and a more rapid rate of cognitive decline. This was followed by Mic.12 (APOE^+^) which was associated with A*β* burden, tau burden but not with cognitive decline. We also found a strong association between the stress response Ast.10 subpopulation, which was present in greater proportion in individuals with greater tau burden and among those with more rapid cognitive decline. Further, we note an association between oligodendrocyte Oli.6 proportion and cognitive decline. Finally, we found two inhibitory neuronal subpopulations associated with tau burden: a positive association for PV+ Inh.16 and a negative association for SST+ Inh.6 (**Fig. 3b**), suggesting relative resilience of PV+ neurons and enhanced vulnerability of SST+ neurons in AD relative to all inhibitory neurons, consistent with prior reports^4, 25^.

**Figure 3:**
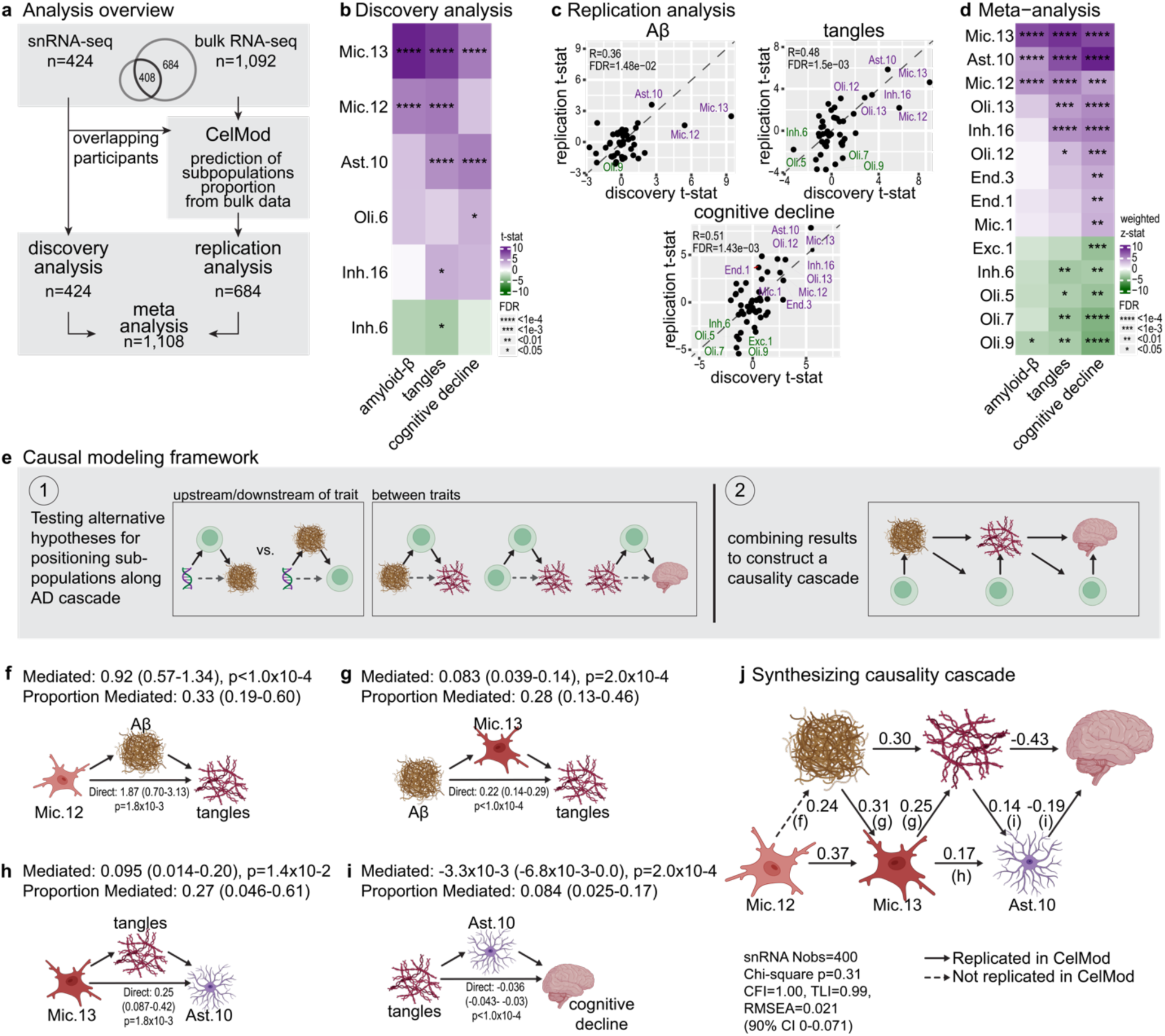
Associating subpopulations with AD-related endophenotypes and positioning them along the AD cascade. (**a**) Overview of our strategy for association analyses with the three prioritized AD-related traits: amyloid b (A*β*), tau tangles, and minus the slope of cognitive decline. The *Discovery* analysis is conducted with 424 participants with snRNA-seq data; the *Replication* analysis is performed using 684 participants with inferred subpopulations proportions from bulk RNAseq data; and a meta-analysis of both sets of data provided a summary of the statistical evidence available to date. (**b**) *Discovery analysis* associating subpopulations to AD traits: snRNA-seq subpopulations significantly associated with at least one of the tested AD-related traits (linear regression with covariates FDR<0.05). n=424 participants. Color scale: association effect size indicating direction and strength from negative (green) to positive (purple) associations. (**c**) *Replication analysis* in an independent cohort aligns with the discovery analysis: Correlation between the *Discovery* (snRNA-seq, n=424, x-axis) and *Replication* (bulk inferred proportions, n=684, y-axis) effect sizes of each subpopulation for each of the three AD-related traits (**d**) Meta-analysis of AD-trait associations over the *Discovery* and *Replication* samples showing subpopulations significantly associated with at least one of the tested AD-related traits. Association direction and strength per subpopulation presented as in (b). (**e**) Causal modeling framework. Each subpopulation associated with endophenotypes can be positioned along the accepted AD cascade (Aβ → tau → cognitive decline) by testing its effect upstream of each endophenotype, mediating between two endophenotypes, or mediating between a different subpopulation and an endophenotype. (**f-i**) Causal mediation models positioning Mic.12, Mic.13 and Ast.10 within the AD cascade. **(f)** A*β* partially but significantly mediates the Mic.12-tau association, placing Mic.12 upstream of A*β*. (g) Mic.13 significantly mediates Aβ-tau association, suggesting that it is upstream of Mic.13. **(h)** Tau partially mediates Mic.13-Ast.10 association, with a substantial direct association between Mic.13-Ast.10. **(i)** Ast.10 partially but significantly mediates the tau-cognitive decline association. (**j**) Structural equation model (SEM) positioning Mic.12, Mic.13 and Ast.10 within the AD cascade. Integration of all the independent mediation results (from f-i and **Supplementary Fig. 5c-f**) in a final structural equation model (SEM). Arrows: Directionality of the association, with relative strength (numeric value), and the guiding mediation model (letter). Solid arrow: the association was replicated (p<0.05) in the Replication sample (n=521 bulk-estimated, with non-missing data for all tested variables, **Supplementary Fig. 5g**). Dashed arrows: The associations was not replicated (p>0.05). CFI=comparative fit index, Nobs=number of observations (participants), RMSEA=root mean square error of approximation, TLI=Tucker Lewis index.

To validate our findings in a larger and independent set of individuals, we used the recently developed CelMod method^4^, to infer the proportion of 92 cell subpopulations in the DLPFC of 1,092 ROSMAP participants who have bulk RNA-seq (excluding rare immune populations, **Methods, Fig. 3a**). We fitted CelMod using profiles of 408 participants for whom we had both snRNA-seq and bulk RNA-seq, and used the proportions of the 45 subpopulations with high confidence predictions for downstream analyses (FDR<0.05, **Methods, Supplementary Fig. 5a**). Using these inferred cell proportions from an independent set of 684 ROSMAP participants (referred to as the *Replication sample*), we replicated our top results and found high correlations between the estimated effect sizes over all subpopulations tested in both the *Discovery*-(n=424 snRNA-seq) and *Replication-* (n=684) *analyses* for all three traits (**Fig. 3c, Supplementary Fig. 5b**).

Finally, to maximize statistical power, we performed a meta-analysis on the full set of 1,108 distinct participants (n=424 discovery and n=684 replication, **Fig. 3a, Methods, Supplementary Table 3**). With the increased statistical power, lipid-associated Mic.12, Mic.13 and stress response Ast.10 had stronger evidence of association with all three traits. The evidence for PV+ Inh.16 and SST+ Inh.6 neurons with tau burden and rate of cognitive decline is also enhanced. The meta-analysis also identified multiple additional subpopulations to be significantly associated with one or more traits, particularly oligodendrocyte and endothelial cell subpopulations (**Fig. 3d**). We thus obtained a robust set of results prioritizing specific neocortical cell subpopulations across different cell types in relation to AD.

### Causal modeling aligns glial subpopulations along the cascade of AD pathophysiology

Next, we sought to test for putative causal relationships relating the proportion of specific subpopulations and AD endophenotypes along the AD pathogenic cascade. To derive a plausible causal sequence, we applied mediation modeling, leveraging the widely accepted cascade of AD progression: Aβ accumulation, followed by tau neurofibrillary tangle formation and finally cognitive decline^26–28^, with the strong genetic risk factor, *APOEε4*, as an anchor for the sequence of events. We first performed causal mediation analysis to quantify direct and mediated (indirect) effects among subpopulations and AD endophenotypes, followed by structural equation modeling to align subpopulations with each step of the AD cascade (**Methods**). We focused on the subpopulations that were most robustly associated with the endophenotypes (association FDR<0.01 with two or more AD-related traits): lipid-associated Mic.12, Mic.13 and stress response Ast.10.

Our model placed Mic.12, Mic.13 and Ast.10 subpopulations at specific points along the AD cascade inferring their putative causal effect: (1) Mic.12 upstream of Aβ (**Fig. 3f, Supplementary Fig 5c**); (2) Mic.13 downstream of Aβ and Mic.12 but predicted to have a direct causal effect upstream of tau and cognitive decline (**Fig. 3g, Supplementary Fig. 5d-e**); and (3) Ast.10 downstream of tau and Mic.13, but upstream of cognitive decline (**Fig. 3h-i, Supplementary Fig. 5f**). We found that Mic.12 had a strong association with age (p=3.2×10^-5^) and thus might drive an age-dependent microglial dysfunction leading to poor Aβ clearance, given its upstream position in the model. Mic.13 proportions mediated 28% of the strong association between Aβ and tau (**Fig. 3g**), yet the strong association between Mic.13 and cognitive decline was mostly mediated through tau (63%). This suggests that the strong influence of Mic.13 on cognitive decline occurs mostly through tau-dependent mechanisms (**Supplementary Fig. 5g**). Unlike Mic.12, Mic.13 was not associated with age but instead was associated with the genetic risk factor *APOEε4*. Further, there was an association between Mic.13 and Ast.10 even after adjusting for tau, suggesting that these two populations may interact separate from tau-related processes (**Fig. 3h**). Finally, Ast.10 mediated 8.4% of the strong tau–cognition association (**Fig. 3i**), and additionally explained 2.9% of the variance in cognitive decline not related to AD pathology. This suggests that Ast.10 might serve as a converging point of tau-dependent and -independent mechanisms leading to cognitive decline.

Synthesizing the above findings, we constructed a structural equation model (SEM) integrating Mic.12, Mic.13, Ast.10, and AD endophenotypes into a single model of the AD cascade (CFI=1.00, TLI=0.99, **Methods, Fig. 3j**). We validated the SEM using the replication sample and reproduced all of the relationships outlined in our snRNA-seq model except the link between Mic.12 and Aβ (**Supplementary Fig. 5g**), demonstrating the robustness of our results and indicating that our modeling strategy has derived plausible causal pathways aligning specific cell subpopulations with key AD endophenotypes, providing a strong set of hypotheses for future mechanistic testing.

### Two distinct disease-associated microglial subpopulations

Our model proposes that Mic.12 and Mic.13 are most likely to play early key causal roles in the proteinopathies of AD. Compared to other microglial cells, both subpopulations have a higher expression of AD-risk genes and share multiple molecular pathways, such as expressing genes involved in foam-cell differentiation, cholesterol storage and lipid metabolic/catabolic process, and lower expression of genes associated with glial cell migration (**Fig. 4a**). However, these two microglial subpopulations also have distinct gene expression patterns. Mic.12 has higher expression of Major Histocompatibility Complex (MHC) class II genes, while Mic.13 upregulated pathways related to negative regulation of locomotion and ECM organization (e.g. ADAM10 and TGFBR1) as well as genes for negative regulation of immune system processes (**Fig. 4b-c, Supplementary Fig. 6a, Supplementary Table 2**). Mic.13 also has higher expression of TREM2, which harbors multiple AD susceptibility variants (**Fig. 4c**).

**Figure 4:**
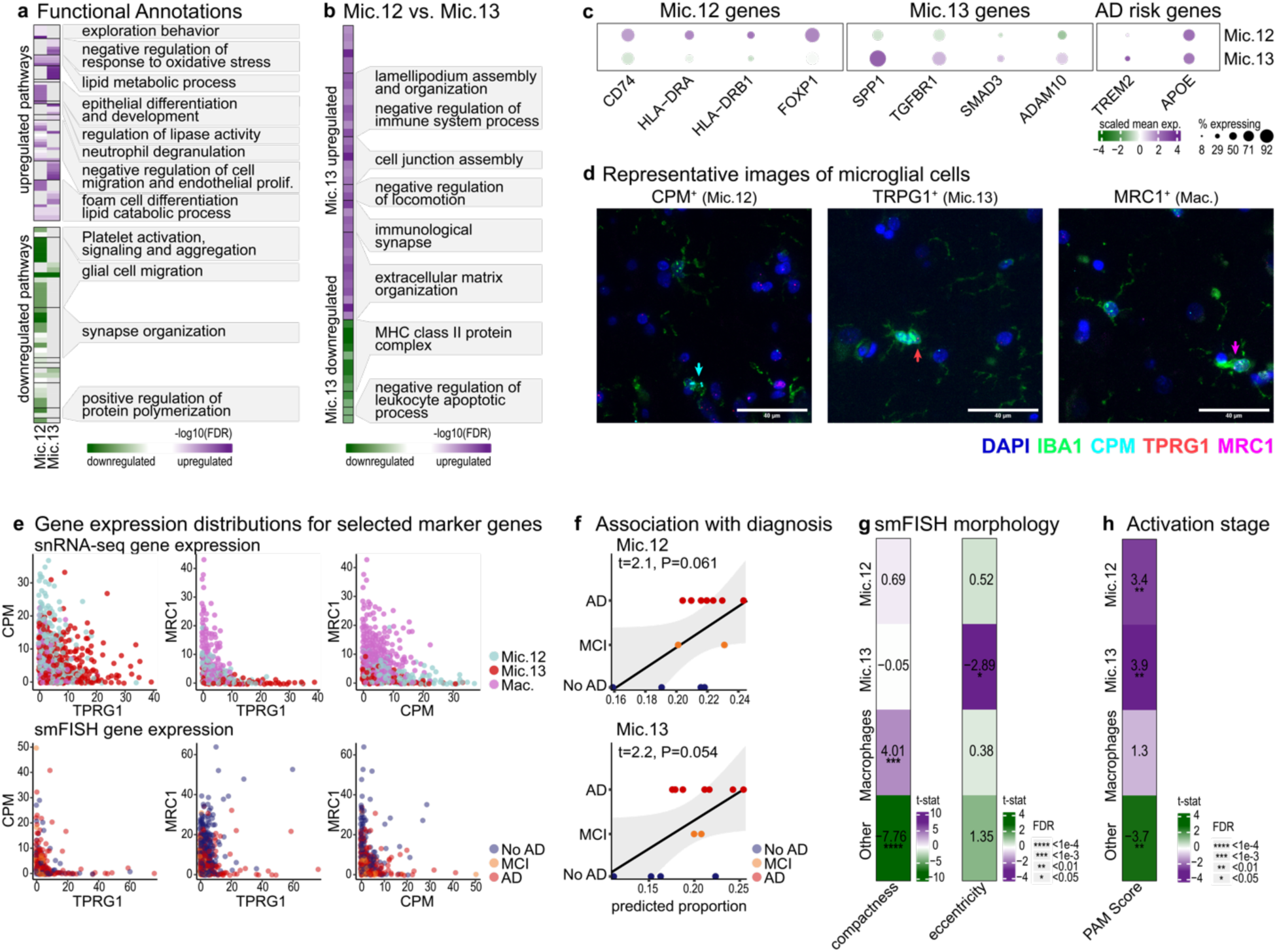
Shared and distinct features of Mic.12 and Mic.13 subpopulations. (**a-b**) Shared and distinct enriched pathways by Mic.12 and Mic.13 subpopulations. Upregulated (purple) or downregulated (green) pathways of: (a) Mic.12 and Mic.13 compared to all other microglial subpopulations, or (b) Mic.13 compared to Mic.12. (**b**). Pathways are grouped based on gene similarities and annotated by representative pathways (**Methods**). Color-scale: −log10 FDR of enrichment (hypergeometric test, p<0.05, **Methods**). (**c**) Unique gene expression pattern for Mic.12 and Mic.13. Gene expression (columns) across Mic.12 an Mic.13 subpopulations (rows) for selected differential genes of interest. Dots color: the mean expression level in expressing cells (column scaled), and dots size: the percent of cells expressing the gene. Full dotplot showing all microglial subpopulations in **Supplementary Fig. 6a**. (**d**) Validation distinct microglial cells expressing the Mic.12 and Mic.13 markers. Representative images from the DLPFC on independent samples (New York Brain Bank) showing: all microglia/myeloid cells (green, anti-IBA1 immunofluorescence), all nuclei (DAPI), RNA scope probes targeting *CPM* (cyan, Mic.12 marker), *TPRG1* (red, Mic.13 marker), and *MRC1* (magenta, macrophage marker). Arrows showing examples of (from left to right) *CPM*^high^, *TPRG1*^high^ and MRC1^high^ cells positive to a single marker. (**e**) Bivariate expression distributions of the three marker genes measured by RNAscope matches snRNAseq. Top row: Expression levels of pairs of marker genes per nuclei (dots) in snRNA-seq data, colored by the atlas annotation (Mic.12, Mic.13 and Macrophage). Bottom row: Expression of pairs of marker genes per myeloid cell (IBA1+) measured by RNAscope data in DLPFC samples of 13 brains from the New York Brain Bank (7 AD, 2 MCI and 4 non-impaired individuals). In each row, the first graph compares the expression of CPM and TPRG1, which shows the bivariate distribution of these markers among microglia: Mic.12 and Mic.13 are at the extreme of each distribution. The second and third graphs compare these microglial markers to the macrophage marker, illustrating they are distinct. (**f**) Association of proportions of Mic.12 (top) and Mic.13 (bottom) microglia from the RNAscope data with the clinical diagnosis of each brain (associations of macrophages and remaining microglial states are shown in **Supplementary Fig. 7c**). Line = linear regression, t=t-statistic, p=p-value. (**g**) Mic.12 and Mic.13 have distinct morphological traits. Associations of Mic.12 and Mic.13 proportions to morphological traits captured by automated segmentation of smFISH images using CellProfiler by IBA1 immunofluorescence to capture the cell morphology (**Methods**). Color scale showing two measures of roundedness: compactness (a measure of how irregular the surface is), and eccentricity (a measure of the extent to which a shape is ovoid). Showing negative eccentricity in Mic.13. (**h**) Association of Mic.13 and Mic.12 proportions to the previous reported activated microglia (PAM) score. PAM score = square root of stage III activated macrophage-appearance microglia density proportion, derived from histological analyses of 92 ROSMAP participants from our Discovery sample; Color scale = association of snRNA-seq proportions to PAM score (histopathology).

To validate and further investigate the distinct features of the Mic.12 and Mic.13 subpopulations, we performed single-molecule RNA FISH (smFISH) experiments in an independent sample of 13 brains from the New York Brain Bank (4 cognitively unimpaired, no AD; 2 MCI; 7 AD). We quantified RNA expression of markers for Mic.12 (CPM), Mic.13 (TPRG1) and macrophages (MRC1), along with immunohistochemical stainings for the general microglial marker IBA1 labeling all myeloid cells (**Fig. 4d, Supplementary Fig. 6b-c**). Out of 16,344 DAPI^+^IBA1^+^ cells, we identified 6,115 cells positive for at least one of the three markers (**Methods**), with smFISH distribution of gene expression levels matching the snRNA-seq counts (**Fig. 4e, Supplementary Table 4**). Matching our clustering analysis, we found that microglia expressing high levels of the Mic.12 marker CPM are distinct from those with high expression of the Mic.13 marker TPRG1, validating that Mic.12 and Mic.13 are indeed distinct microglial populations (**Fig. 4d-e**).

Next, we validated the association of Mic.12 and Mic.13 proportions with AD diagnosis. We quantified the smFISH proportions for each of Mic.12, Mic.13 and macrophages subpopulations (**Methods, Supplementary Fig. 6d-e**). Despite the limited number of samples and the inherently large variability of the data, we found trend level associations between the proportions of Mic.12 and Mic.13, which were on average higher in individuals diagnosed with MCI, and even more so in AD, as compared to cognitively unimpaired (**Fig. 4f**).

Finally, we tested whether Mic.12 and Mic.13 cells are morphologically distinct. We observed a specific morphological identity to the Mic.13 subpopulation, with significantly lower eccentricity (compared to all other microglia, **Methods, Fig. 5g**). To further explore the morphological features of these subpopulations, we leveraged morphology-based quantification of microglial activation stages (stages I-III) in the midfrontal cortex, available for 92 participants within our snRNA-seq cohort. Stage III microglia, are activated macrophage-appearance microglia, were shown to be more condensed and less ramified, and were shown to increase in proportion with AD and its endophenotypes^29^. We calculated the Proportion of Activated Microglia (PAM) score^29^ per individual, which is the square-root of the proportion of stage III microglia. We found that Mic.12 and Mic.13 have a strong positive association to the PAM score, while all other microglia have a negative association (**Fig. 4h**). Of note, Mic.12 association with PAM is attenuated when adjusted for Mic.13 suggesting that Mic.13 may be the primary driver of this association (**Supplementary Table 4**).

**Figure 5:**
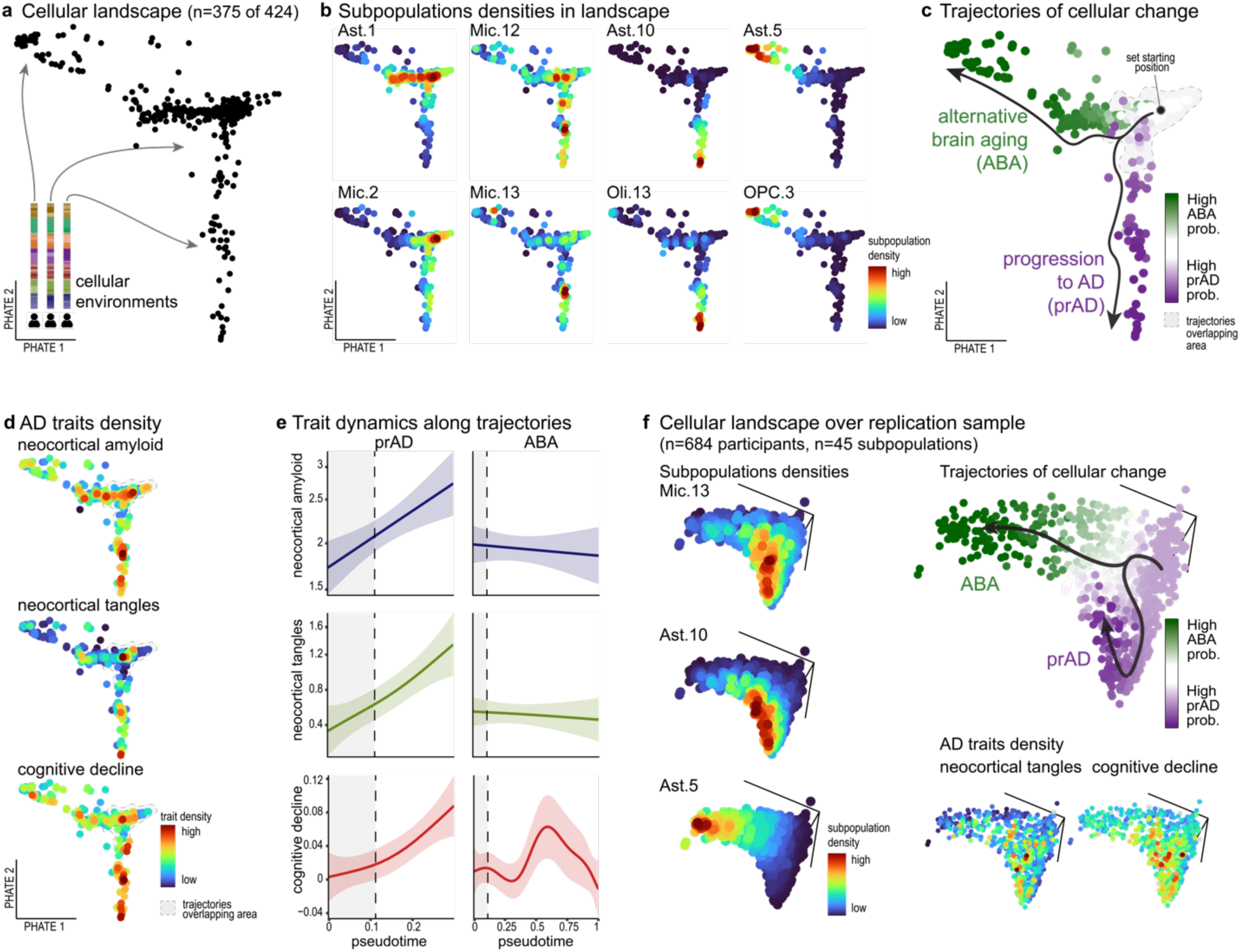
Modeling the cellular landscape manifold uncovered two distinct trajectories of brain aging, one leading to AD dementia. (**a**) The structure of the cellular landscape manifold of the aged neocortex. The cellular manifold captured by BEYOND algorithm for the discovery sample (n=424). Showing PHATE^56^ embedding of each participant (individual dots) based on similarity of their cellular environments, represented as the cellular composition profiles in our cell atlas of 92 subpopulations (**Methods**). See also **Supplementary Fig. 7a** (**b**) Distinct patterns of subpopulation proportions along the cellular landscape manifold. Participants are colored by the locally-smoothed proportions of each subpopulation (embedded in the cellular landscape as in a), showing distinct patterns for different subpopulation for representative examples. (**c**) Two uncovered cellular trajectories of brain aging. Two predicted trajectories in the cellular landscape manifold, termed progression of AD (prAD) and Alternative Brain Aging (ABA). Cellular landscape (as presented in a) color-coded by the difference of the participant’s probability of belonging to the prAD trajectory and the ABA trajectory. The area with high overlap between the trajectories (see **Supplementary Fig. 7d**) is marked in grey. Arrows illustrate the pseudotime inferred directionality of the trajectories starting from the shared root participant and going towards the terminal states (see pseudotime color-coding in **Supplementary Fig. 7c**). (**d-e**) Distinct patterns and dynamics of AD-traits along the prDA and ABA trajectories. Description of (top) neocortical A*β*, (center) neocortical tau and (bottom) cognitive decline. Showing: (**d**) distinct patterns in the cellular landscape manifold per trait. Participants are color-coded by the locally-smoothed density values of the traits in the landscape (as in a); and (**e**) Distinct trait dynamics in each trajectory. Trait-dynamics along the pseudotime in each of the inferred trajectories. (**f**) Validation of the cellular landscape manifold and the trajectory analysis over an independent cohort. Applying BEYOND to the Replication sample (n=684) using only the 45 well-predicted subpopulations (**Supplementary Fig. 5a**). Showing for the replication manifold - Left: Distinct densities (as in b) for key subpopulations; Top-right: Two inferred trajectories of cellular changes (as in c); and Bototm-right: Distinct AD-trait densities separate into two directions, similar to the patterns over the discovery manifold.

### Reconstruction of cellular trajectories uncovered two paths that reflect distinct disease states

Although the association and causality analyses highlight key subpopulations and predict their causal role along the disease cascade, they do not provide a complete view of the cellular-molecular changes underlying AD progression, especially in its early stages, and of how it might differ from natural brain aging. Such analysis, however, is limited by the lack of longitudinal molecular brain measurements, as specimens are obtained at autopsy and miss temporal cellular dynamics within each individual. For dynamic analysis, all participants need to be aligned along a timeline based on their disease stage, which we cannot accurately infer based on their clinicopathologic profiles alone. We thus devised an alternative approach, using a new conceptual-analytical framework we call BEYOND (**Fig. 1a**): an integrative approach to uncover one or more trajectories of cellular changes, map them to clinicopathological outcomes and accurately align participants along them. BEYOND represents each participant by their cellular composition profile in our cell atlas to model their *cellular environment*, and builds a *cellular landscape* manifold that captures the diversity of cellular environments, with each participant as a single point in the manifold (**Fig. 5a-b**). It then reconstructs trajectories of change along the manifold using similarities in the cellular environments across participants (**Fig. 5c**). Next, it assesses whether the trajectories are associated with AD-related traits, matching them to potential disease states (**Fig. 5d-f**). Finally, BEYOND assigns cell subpopulations to multi-cellular communities by integrative clustering of subpopulations with similar proportions across individuals and dynamics along the inferred trajectories (**Fig. 6a, Methods**).

**Figure 6:**
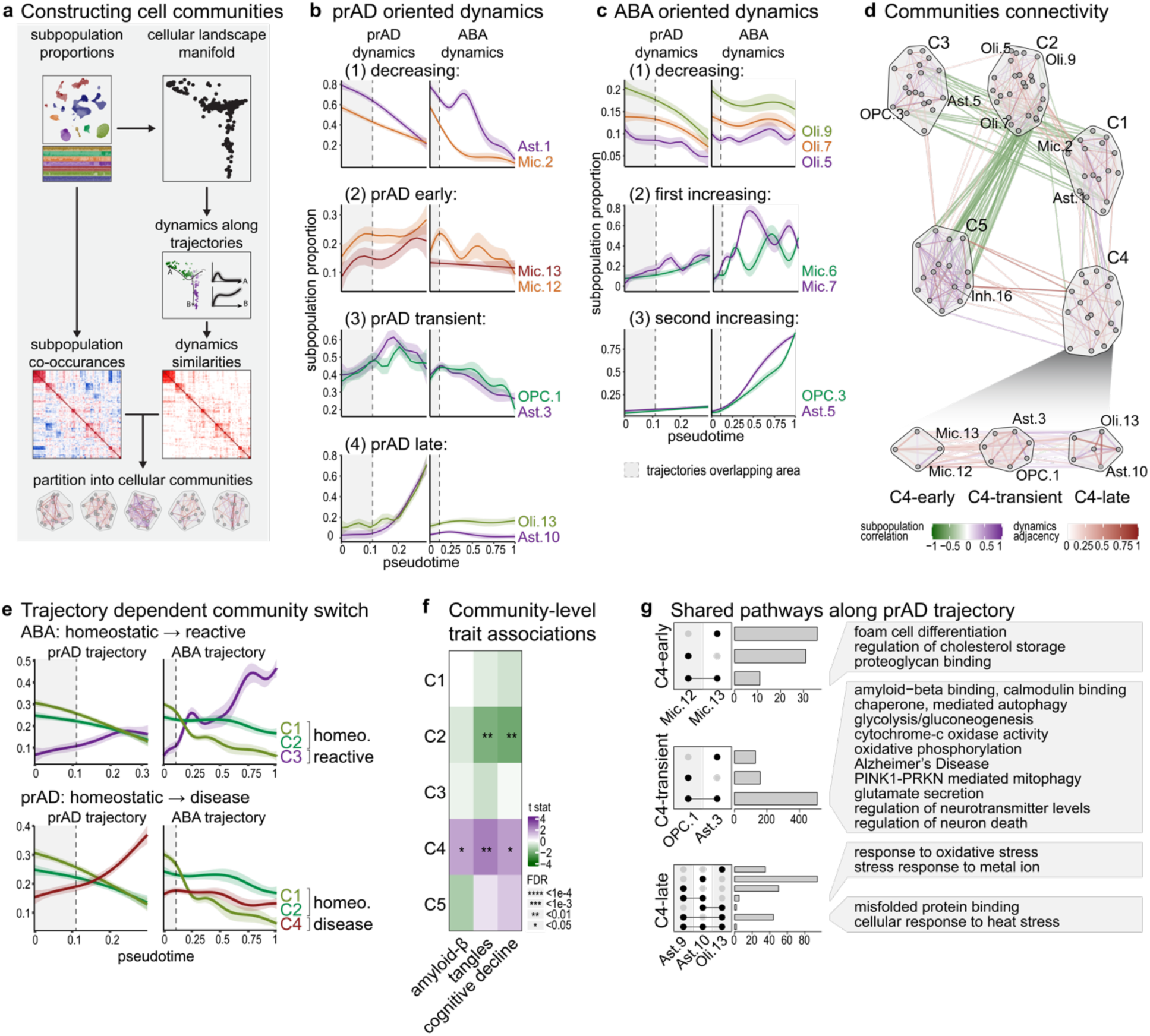
Multicellular communities with distinct dynamics along the prAD and ABA trajectories. (**a**) Schematics of the BEYOND algorithms for construction of cellular communities based on both subpopulation dynamics and -co-occurrences across individuals. (**b,c**) Distinct dynamical patterns of subpopulations across trajectories. Subpopulation with trajectory-specific dynamics assigned to the (b) prAD trajectory and (c) alternative-aging trajectory. Trait-dynamics along the pseudotime in each of the inferred trajectories, prAD (left) and ABA (right), uncovering subpopulations with early, transient and late patterns. (**d**) Multi-cellular communities of cell subpopulations with shared dynamic patterns and co-occurrences. Subpopulation node graph with subpopulations as nodes and similarity between subpopulations as edges. Green-purple edges are colored by co-occurrences (green for negative and purple for positive), red edges are colored by dynamics similarity. Communities C1-C5 are marked on the graph. Bottom inset: Node graph as above showing sub-clustering of disease community C4 to three sub-communities: early-C4, transient-C4 and late-C4. Edges of correlation between(−0.2, +0.4) or dynamics similarity between (−0.2, +0.2) were excluded from visualization. Full connectivity is shown in **Supplementary Fig. 8c**. (**e**) Distinct dynamic patterns for cellular communities across trajectories. Whole-community dynamics showing change in community proportion (**Methods**) along pseudotime in the two trajectories. (**f**) Associations of cellular communities C2 and C4 to AD-traits. Trait associations showing the effect size of a community’s proportion per individual on AD endophenotypes (linear regression T-test, FDR<0.05). (**g**) Shared and distinct pathways found for main subpopulations within the the C4-early, -transient and -late sub-communities.

When applying BEYOND to the cellular composition profiles in our cell atlas (**Fig. 1h**) and investigating the manifold in a two-dimensional space, we noticed two major axes of change (**Fig. 5a, Supplementary Fig. 7a**). Along each axis we found distinct patterns of multiple subpopulations across all cell types, indicating that the trajectories indeed captured changes in the overall cellular environment (**Fig. 5b**). For example, the disease associated subpopulations Mic.12, Mic.13 and Ast.10 all increase in proportion along one axis, while other subpopulations of reactive glial cells such as Ast.5 and OPC.3 (SERPINA3^+^OSMR^+^) increase along the other axis. These divergent cellular patterns are robust to both the embedding algorithm and to the number of cell subpopulations used as input (**Supplementary Fig. 7b**).

We used the *cellular landscape* manifold to uncover trajectories of cellular change and assign a pseudo-time ordering of the participants along these trajectories (by *Palantir*^30^, **Methods**). We inferred two divergent trajectories from a shared starting point in the manifold (**Fig. 5c, Supplementary Fig. 7c-d**). The trajectory analysis is blind to the participants’ clinicopathological characteristics, enabling association of AD traits to trajectories (**Methods, Fig. 5d-e, Supplementary Fig. 7e**). The resulting analysis suggests that one of the two trajectories is strongly related to AD, which we named “progression to AD” (prAD) that captured a monotonous increase in neocortical A*β* and tau burden as well as an increase in the rate of cognitive decline; We named the other trajectory “Alternative Brain Aging” (ABA), as it captured individuals with a constant level of A*β*, low/no neocortical tau, and a slower rate of cognitive decline with a variable pattern. The variability in the pattern of decline along this second trajectory may indicate that it is influenced by comorbidities, other neuropathologies contributing to cognitive decline and/or resistance mechanisms that are yet unappreciated.

We validated our discovery of the two distinct trajectories that capture different disease courses, by applying the BEYOND methodology to the replication sample of 684 additional participants (with CelMod estimated proportions for 45 subpopulations, **Methods, Fig. 3a**). We recapitulated the two main trajectories associated with divergent cellular environments and AD-related traits, matching the prAD and ABA trajectories (**Fig. 5f, Supplementary Fig. 7f-h**). Our ability to replicate these findings in an independent set of participants, with only a partial set of subpopulations, highlights the robustness of our findings. BEYOND was thus able to reconstruct a timeline that captures the continuous changes along a trajectory leading to AD, and second major trajectory of alternative brain aging, while only using the cellular architecture of the DLPFC in each participant. Interestingly, individuals along the ABA trajectory are not completely free from cognitive decline, but the difference in the cellular cascades between the prAD and ABA trajectories suggests that they will each need to be considered separately in therapeutic trials.

### Diverse multi-cellular communities with distinct dynamics along disease and aging timelines

To gain a deeper understanding of the changes in the cellular environment of the brain underlying the progression of AD and alternative brain aging, we examined the coordinated dynamics of all cell subpopulations along the two discovered trajectories (**Fig. 6a**). The alignment of participants along a timeline by BEYOND enabled the inference of dynamics of each subpopulation along the prAD and ABA trajectories (**Methods**). At the individual subpopulation level, we found different patterns of subpopulation dynamics, including: prAD-specific subpopulations and ABA-specific subpopulations that changed in proportion along one trajectory but not the other; or temporal-specific subpopulations along the prAD trajectory, differentiating between early, late or transient patterns (**Fig. 6b-c, Supplementary Fig. 8b**).

To find multicellular communitie*s*^4^ that reflect coordinated changes within cellular environments, we applied an integrative clustering step (the final step in the BEYOND methodology, **Fig. 6a**). We clustered subpopulations using two properties reflecting different aspects of coordinated change: (1) *co-occurrences* of subpopulations across individuals (similar to ^4^) and (2) shared patterns of subpopulation dynamics along all trajectories (**Methods**). The clustering revealed five cellular communities - which we named C1-C5 (**Fig. 6d, Supplementary Fig. 8c**). Within each community, subpopulations were found to be highly inter-connected, with similar occurrences and dynamics, while between communities the degree of similarity varied and was dependent on their proximity within the cellular manifold (**Supplementary Fig. 8d**).

Communities C1 and C2 were composed of subpopulations decreasing in proportion along both trajectories (**Fig. 6b-c**). The C1 community included homeostatic microglial and astrocytic subpopulations (e.g. Ast.1, Mic.2), which decreased in proportion faster towards the ABA trajectory, while the C2 community included SST+ neurons (Inh.6) and oligodendrocyte subpopulations (Oli.5, Oli.7, Oli.9) negatively associated with cognitive decline, which decreased in proportion faster towards the prAD trajectory compared to the ABA trajectory (**Fig. 3d, Fig. 6b-c,e**). The ABA specific community C3 included the reactive-like subpopulations of microglial (Mic.6-7), astrocytes (SERPINA3^+^OSMR^+^ Ast.5) and OPCS (SERPINA3^+^OSMR^+^ OPC.3, **Fig. 3d, Fig. 6c, Supplementary Fig. 8b**). The neuronal community C5 included PV+ Inh.16 which was found to be negatively associated with tau burden, and had minimal change in proportions mainly along the prAD trajectory (**Fig. 3b, Supplementary Fig. 8b,d**). Finally, we found a prAD specific community C4, which included subpopulations that increased in proportion along the prAD trajectory. The AD-associated glial subpopulations (by trait association analysis, **Fig. 3b,d**), Mic.12, Mic.13, Ast.10 and Oli.13 are all found to be part of the prAD community C4, and their different dynamic profiles align with our causal modeling results (**Fig. 3f-j**). We next associated AD-traits to each community, and found that C4 proportions are positively associated with neocortical A*β* and tau as well as cognitive decline, while increased proportions of the cognitive-healthy community C2 negatively associated with neocortical tau and cognitive decline (**Methods, Fig. 6f**).

To better explore the dynamics along the prAD trajectory, we further sub-clustered the prAD specific community C4, which was divided into three multicellular sub-communities(**Fig. 6d, Methods**): (1) C4-early - consisting of cell subpopulations increasing early in prAD, including the AD-associated Mic.12 and Mic.13, (2) C4-transient - consisting of subpopulations increasing transiently along the trajectory, such as enhanced-mitophagy subpopulations Ast.3 and OPC.1, and (3) C4-late - consisting of subpopulations increasing late along the prAD, including the stress response and AD associated Ast.10 and Oli.13 subpopulations. Of note, we found high coordination within the co-occurrence of all subpopulations in the prAD community C4, yet their dynamic profiles distinguish these three C4 sub-communities (**Supplementary Fig. 8c**). Thus, the dynamic reconstruction expands our perspective beyond the trait-association analysis (**Fig. 3**), highlighting the transitional state before terminal AD that is defined by specific multicellular communities.

To explore changes in molecular processes along the progression of AD, we next searched for shared functionality within each of the three prAD-specific sub-communities, the early-C4, transient-C4 and late-C4, to align specific pathways to each disease stage. Within the C4-early community lipid metabolism and immune related pathways were shared between Mic.13 and Mic.12 (as described above, **Fig. 4**). Within the C4-transient community, Ast.3 and OPC.1 both upregulated mitochondrial and mitophagy pathways, metal-ion uptake and glucose metabolism genes as well as upregulated genes linked to neurotransmitters and glutamate secretion. Finally, within the C4-late community, Oli.13, Ast.9, Ast.10, End.5 and Mic.11, all upregulated a diversity of stress response pathways, specifically sharing the upregulation of genes linked to unfolded protein response, ROS, oxidative stress and metal-ion stress (**Fig. 6g**). Thus, we see a progression of molecular pathways being engaged in the prAD trajectory, starting with lipid-/inflammation-related ones early on that are followed by transient changes in glucose metabolism/mitochondrial-/synaptic-function and end with a broader metabolic shift involving reactive oxygen species and stress responses at late stages.

The finding of distinct multicellular communities with correlated proportions across participants and similar dynamics along both trajectories seen among aging brains, highlights the coordinated changes in cellular populations that underlie both progression to AD and the ABA trajectory, that remains less well defined. These results confirm a cellular cascade and molecular dynamics that are specific for AD and appear to be absent from a large portion of aging individuals which are following the alternative path of brain aging.

## Discussion

Here, we uncovered the cellular landscape and dynamics of the aged human brain, defined using a high resolution cell atlas from snRNA-seq profiling of a large sample of 465 ROSMAP participants with deep clinical- and neuropathological phenotyping (424 with sufficient number of nuclei for the downstream analysis). The combination of our unique set of participants, detailed atlas and newly devised algorithmic approach BEYOND, allowed us to provide insights that address fundamental open questions regarding the process of aging and AD in human brains: **First**, on the question of understanding the inter-individual heterogeneity seen in the older brains, we showed the presence of two distinct cellular trajectories, distinguishing AD from alternative brain aging already at the earliest stages of the disease. **Second**, we report the importance of cellular changes in defining these trajectories: reconstructing pseudo-temporal cellular dynamics, uncovered cellular changes unique to each of the trajectories, distinguishing between early, transient and late changes. **Third**, individual subpopulations do not contribute by themselves: we uncovered archetypical multicellular communities, whose subpopulations share unique dynamics and molecular pathways, facilitating the coordinated shift in cellular environments underlying variations in the cellular landscape. **Fourth**, our analyses highlight communities and key subpopulations contributing to AD and also proposed where, in the cascade of events linking amyloid- and tau proteinopathies to cognitive decline they may exert their casual role. Thus, we contribute insights into both the conceptual framework underlying AD therapeutic development by highlighting the need to engage with specific cellular communities at different points along the causal chain leading to AD and to the trial design that needs to recognize and exclude the subset of individuals that are on the alternative brain aging path that are not likely to respond to interventions geared at the cellular changes of the trajectory to AD.

Our results and insights rest on a foundation of high-quality snRNA-seq data generation coupled to the deployment of innovative analytic approaches and use of rigorous statistical methods. Specifically, we provide evidence of replication for both our results at the level of associations of individual cell subpopulations to AD traits and the results of our modeling that leads to our trajectories of brain aging and cellular communities in an independent cohort of 684 participants. This is thanks to the CelMod algorithm^4^ that enables us to infer subpopulation proportions from bulk RNA-seq data. All snRNA-seq data, inferred proportions from bulk data and analytic results have been uploaded to the AD Knowledge Portal to facilitate repurposing. As one of the largest single nucleus studies performed to date, we also offer important insights for future study design and power calculations given that we have successfully identified and replicated associations with multiple different cell subpopulations: with our n=424 participants, we were adequately powered for the analyses we conducted.

Our work makes several significant advances in our understanding of the sequence of events leading to AD and brain aging, showing that there are continuous, stereotypic, reproducible changes in cellular environments throughout the causal chain leading to AD and different changes leading to an alternative form of brain aging (**Fig. 6**). Previous snRNA-seq studies have associated cells and pathways to AD^4–15^, but were limited to a case-control study design and thus could neither infer cellular dynamics nor decouple AD and alternative aging. Because we use a sample of the older population and a prospective cohort study design, we leverage the full diversity of the older brain, allowing us to reconstitute the different trajectories of brain aging in a data-driven manner by applying BEYOND. Specifically, we suggest the following sequence of events underlying the prAD trajectory: **At the early stages**, selective homeostatic glial subpopulations decrease in proportion alongside an increase of, first, the lipid-associated microglia subpopulation APOE^+^ Mic.12 subpopulation that is itself influenced by advancing age and contributes to the accumulation of A*β* proteinopathy, and up-regulates immune activation pathways. Then, a distinct but related Mic.13 subpopulation of APOE^+^TREM2^+^ microglia that are influenced by APOEε4 (the strongest genetic risk factor for AD) contributes to the subsequent accumulation of tau proteinopathy (**Fig. 3**). **At the next stage** of the AD cascade, with the accumulation of tau proteinopathy, we observed a transient increase in the proportions of Ast.3 and OPC.1, which both upregulate genes associated with high energy demand and enhanced-mitophagy, as well as oxidative phosphorylation and glutamate secretion (**Fig. 7**). OPC.1 further upregulates genes associated with response to oxidative stress aligning with reports suggesting the increased vulnerability of OPCs^31^ to oxidative stressors^31^ that are rising during this phase. **At the last stage** of the AD cascade, we observed further increase in Mic.12 and Mic.13 proportions, together with a coordinated increase of Ast.10 and Oli.13, with Ast.10 playing an important role mediating the effect of tau proteinopathy on the increased rate of cognitive decline. Both Ast.10 and Oli.13 express stress response genes, with Ast.10 mainly demonstrating response to oxidative stress while Oli.13 showing response to heat stress and unfolded protein. Cognitive decline appears to be directly affected by Ast.10 that is driven by both tau and Mic.13, suggesting that the proportion of this astrocyte subpopulation may be a point of convergence for different processes leading to cognitive dysfunction. While Mic.12 offers a good target with which to perturb the accumulation of A*β* proteinopathy to enhance therapeutic options centered on anti-A*β* antibodies, preventing polarization of microglia and astrocytes into Mic.13 and Ast.10 respectively, may have more immediate impact in helping to prevent cognitive impairment. The latter strategy would also be better suited for individuals who are already A*β*+ and are at risk of tauopathy. On the other hand, these subpopulations do not appear to be relevant to the **alternative brain aging** (ABA) trajectory, where we found a selective decrease in homeostatic glial subpopulations and an increase in reactive microglial subpopulations. At the next stage, we observed increased proportions of reactive-like Ast.5 and OPC.3, both expressing the markers SERPINA3 and OSMR. Among participants in ABA, we found constant levels of neocortical amyloid, very limited neocortical tau and varying dynamics of cognitive decline. Thus, more work is needed to better understand this trajectory and its interesting, defining glial subpopulations.

Methodologically, our work solved critical challenges in the study of long dynamic processes, as there are potential overlapping molecular processes, such as in AD and brain aging, limiting the ability to position individuals along the stages of the process. First, rigorous statistical methodology, such as mediation analysis, enabled us to propose the most likely scenarios given the limitations of postmortem datasets (**Fig. 3**). Further, we showed that predicting cellular compositions from bulk RNAseq (by CelMod^4^) could be used for validations in a large independent cohort and to increase the sample size in a meta-analysis approach (**Fig. 3, Fig.5**). Finally, the BEYOND strategy enabled the reconstruction of cellular-molecular dynamics linked to disease outcomes: by aligning individuals along a cellular cascade, free of clinical assumptions, we showed that disease timelines as well as unexpected cellular trajectories can be uncovered, and this approach can untangle overlapping processes (**Fig. 5**). Moreover, clustering cell subpopulations enabled the identification of coordinated cellular communities, reflecting changes in entire cellular environments which we suggest are the ultimate therapeutic target (**Fig. 6**). BEYOND and our statistical modeling provide a conceptual framework widely applicable to study any dynamic process with coordinated changes in cellular environments. The BEYOND approach can be generalized to achieve a broader view of trajectories of change in cellular environments, by additional data from genetically diverse individuals, diverse pathological states, multiple brain regions, as well as additional modalities beyond RNA such as proteomics, imaging and chromatin states.

We note that our work has certain limitations. Principally, our sample size was adequate for discovering two trajectories and to prioritize individual cell subpopulations as associated with disease; however, we need larger sample size to elaborate the detailed architecture of the different cellular communities, to better characterize the ABA trajectory, and to perhaps uncover additional trajectories that are not yet apparent. The nature of snRNA-seq data is also a limitation, as we lack the cytoplasmic transcriptome which may harbor distinct pieces of information that may clarify certain points, post-transcriptional modifications and spatial information. Further, while the ROSMAP collections are superb in their design and detailed phenotyping, they are composed largely of individuals of European ancestry, and additional data from other human populations are sorely needed to generalize our results. Finally, we focused on producing data from unsorted nuclei, which preserves the inter-cellular relationships but leads to limited sampling of less common cell subpopulations.

Overall, we present new insights into the cellular and molecular cascades leading to AD and brain aging processes that are based on replicated results from different statistically rigorous methodologies, and prioritized specific pathways and cellular environments as alternative drivers of disease or resilience. Our work highlights that AD is a disease of the whole brain tissue and can’t be effectively approached by engaging any single molecule in a single target cell: we rather need to modulate distinct cellular communities at different stages of AD to restore homeostasis and preserve cognitive function. Further, our data have generated mechanistically precise hypotheses as to where in a sequence of events a particular subpopulation or community may have a role, informing future therapeutic development and clinical trial design. We provided a cellular foundation for a new perspective of AD pathophysiology in which a pathologic cellular community becomes the true target of therapeutic development, and the shared molecular signals of this community provide an important new substrate for the development of new interventions.

## Methods

### Experimental design: Study participants and AD traits

Data were derived from subjects enrolled in one of two longitudinal clinical-pathologic cohort studies of aging and dementia, the Religious Orders Study (ROS) and the Rush Memory and Aging Project (MAP), collectively referred to as ROSMAP. All participants are without known dementia at enrollment, and have annual clinical evaluations and agree in advance to brain donation at death. At death, the brains undergo a quantitative neuropathologic assessment, and the rate of cognitive decline is calculated from longitudinal cognitive measures that include up to 25 yearly evaluations^32^. Each study was approved by an Institutional Review Board of Rush University Medical Center. All participants signed an informed consent, Anatomic Gift Act, and repository consent. For this study, we selected 465 participants, blind to their neuropathologic and clinical traits, and based on availability of frozen pathologic material from the Dorsolateral Prefrontal Cortex (DLPFC), including only participants with RIN>5 and post mortem interval (PMI) <24 hours, as in our prior studies^4^. After defining our cell subpopulations in a subset of 465 participants meeting our quality criteria during data preprocessing, we excluded 19 participants without whole genome sequence data or with missing information, leaving 424 participants retained for disease-association and all other downstream analyses (**Supplementary Fig. 1b**). Our study cohort includes diverse individuals across the full range of the pathological and clinical stages of AD. The demographic and clinicopathologic characteristics are described in **Supplementary Table 1**.

Pathological measures were collected as part of the ROSMAP cohorts (previously described in details^33–35^). We focused our analysis on three quantitative AD related traits, which have a larger statistical power compared to discrete classifications: **Rate of cognitive decline**: Uniform structured clinical evaluations, including a comprehensive cognitive assessment, are administered annually to the ROS and MAP participants. The ROS and MAP methods of assessing cognition have been extensively summarized in previous publications^36–38^. Scores from 19 cognitive performance tests common in both studies, 17 of which were used to obtain a summary measure for global cognition as well as measures for five cognitive domains of episodic memory, visuospatial ability, perceptual speed, semantic memory, and working memory. The summary measure for global cognition is calculated by averaging the standardized scores of the 17 tests, and the summary measure for each domain is calculated similarly by averaging the standardized scores of the tests specific to that domain. To obtain a measurement of cognitive decline, the annual global cognitive scores are modeled longitudinally with a mixed effects model, adjusting for age, sex and education, providing person specific random slopes of decline (which we refer to as ***cognitive decline***). Further details of the statistical methodology have been previously described^39^. **Aβ and tau pathology burden**: Quantification and estimation of the burden of parenchymal deposition of Aβ and the density of abnormally phosphorylated tau-positive neurofibrillary tangles levels present at death (which we refer to as Aβ and tau pathology, respectively). Tissue was dissected from eight regions of the brain: the hippocampus, entorhinal cortex, anterior cingulate cortex, midfrontal cortex, superior frontal cortex, inferior temporal cortex, angular gyrus, and calcarine cortex. 20µm sections from each region were stained with antibodies for the Aβ and tau protein, and quantified with image analysis and stereology. Measurements were summarized to provide a global measure of Aβ and tau burdens. In the analyses of trait associations, causality modeling and dynamics modeling we used the measurements of Aβ and tau midfrontal cortex (referred to as neocortical Aβ and -tau). Further, we note that we are assessing cellular changes in the fresh-frozen DLPFC samples from one hemisphere of each brain in relation to the measures of cortical A*β* and tau burdens measured in the midfrontal cortex of the opposite, fixed hemisphere in which the standard, structured neuropathologic assessment is conducted. Additionally, we make use of quantifications of: final cognitive diagnosis^40^, Braak stage^21, 41^, CERAD score^21, 42^ and NIA-Reagan criteria^21, 22^

### Nucleus isolation and single-nucleus RNA library preparation

To increase library throughput, reduce batch effects and price, we profiled single nuclei in pooled batches of samples. Each batch of samples for library construction consisted of 8 participants, except batch B63 that consisted of 7 participants. Batches were designed to balance clinical and pathological diagnosis and sex as much as possible (**Supplementary Fig. 1a**). DLPFC tissue specimens were received frozen from the Rush Alzheimer’s Disease Center. We observed variability in the morphology of these tissue specimens with differing amounts of gray and white matter and presence of attached meninges. Working on ice throughout, we carefully dissected to remove white matter and meninges, when present. The following steps were also conducted on ice: about 50-100mg of gray matter tissue was transferred into the dounce homogenizer (Sigma Cat No: D8938) with 2mL of NP40 Lysis Buffer [0.1% NP40, 10mM Tris pH 8.0, 146mM NaCl, 1mM CaCl2, 21mM MgCl2, 40U/mL of RNAse inhibitor (Takara: 2313B)]. Tissue was gently dounced 25 times with Pestle A, followed by 25 times with Pestle B while on ice, then transferred to a 15mL conical tube. 3mL of PBS + 0.01% BSA (NEB B9000S) and 40U/mL of RNAse inhibitor were added to a final volume of 5mL and then immediately centrifuged with a swing bucket rotor at 500g for 5 mins at 4°C. Samples were processed 2 at a time, the supernatant was removed, and the pellets were left on ice while processing the remaining tissues to complete a batch of 8 samples. The nuclei pellets were then resuspended in 500ml of PBS + 0.01% BSA and 40U/mL of RNAse inhibitor. Nuclei were filtered through 20µm pre-separation filters (Miltenyi: 130-101-812) and counted using the Nexcelom Cellometer Vision and a 2.5ug/ul API stain at 1:1 dilution with cellometer cell counting chamber (Nexcelom CHT4-SD100-002). 5,000 nuclei from each of 8 participants were then pooled into one sample, and the 40,000 nuclei in around 15-30ul volume were loaded into two channels on the 10X Single Cell RNA-Seq Platform using the Chromium Single Cell 3’ Reagent Kits version 3. Libraries were made following the manufacturer’s protocol. Briefly, single nuclei were partitioned into nanoliter scale Gel Bead-In-Emulsion (GEMs) in the Chromium controller instrument, where cDNAs from the same cell shares a common 10X barcode from the bead. Amplified cDNA is measured by Qubit HS DNA assay (Thermo Fisher Scientific: Q32851) and quality assessed by BioAnalyzer (Agilent: 5067-4626). This WTA (whole transcriptome amplified) material was diluted to <8ng/ml and processed through v3 library construction, and resulting libraries were quantified again by Qubit and BioAnalzyer. Libraries from 4 channels were pooled and sequenced on 1 lane of Illumina HiSeqX by The Broad Institute’s Genomics Platform, for a target coverage of around 1 million reads per channel. The same libraries of batches B10–B63 were resequenced at The New York Genome Center using Illumina NovaSeq 6000. Sequencing data of both Broad Institute and New York Genome Center were used for analysis.

### Pre-processing and quality control steps for snRNA-seq data

For each of the 128 pooled libraries we performed the following steps: (1) library alignment and background noise removal; (2) de-multiplexing; (3) application of a normalization and clustering pipeline; (4) classification of nuclei for cell types; (5) removal of low-quality nuclei with a cell-type specific threshold; (6) detection of doublets for removal. In more details about each step:

#### Library alignment and background noise removal

Libraries were aligned to the GRCh38 pre-mRNA transcriptome and unique molecular identifier (UMI)-collapsing were inferred using the CellRanger^43^ toolkit (version 6.0.0, chemistry V3, 10X Genomics). To control for technical artifacts of background ambient RNA molecules, we ran CellBender^44^ *remove-background* utility over the gene expression matrices generated by CellRanger. Briefly, CellBender is an unsupervised method for inferring empty- and cell-containing droplets, learning the background RNA distribution of the empty droplets, and removal of such technical background noise. We thus retrieve uncontaminated cell-containing droplets (cuda flag set, epochs=300, learning rate=1e-5, z-dim=50). Each of the 128 libraries was processed while setting the number of expected cells according to the number of nuclei estimated by CellRanger.

#### De-multiplexing

We demultiplexed nuclei and inferred participants of origin in our pooled libraries using available genotype data of the participants. Based on participants’ polymorphic sites and each nucleus’ genotype data obtained from the snRNA-seq reads, we assigned each nucleus back to its original participant using the demuxlet software^23^. From the WGS-based VCF file of 1,196 ROS/MAP individuals, we extracted SNPs that were in transcribed regions, passed a filter of GATK, and at least one of the eight individuals had its alternate allele. The extracted SNP genotype data were fed to demuxlet along with a BAM file generated by CellRanger. For libraries in which not all eight individuals had previously been genotyped, we used *freemuxlet* (https://github.com/statgen/popscle), which clusters droplets based on SNPs in snRNA-seq reads and generates a VCF file of snRNAseq-based genotypes of the clusters. The number of clusters was specified to be eight. The snRNA-seq-based VCF file was filtered for genotype quality > 30 and compared with available WGS genotypes using the bcftools *gtcheck* command. Each WGS-genotyped individual was assigned to one of droplet clusters by visually inspecting a heatmap of the number of discordant SNP sites between snRNAseq and WGS. The above two procedures converged to a table that mapped droplet barcodes onto inferred individuals.

#### Normalization and clustering pipeline

The following pipeline was executed on the RNA count matrix: normalization and scaling by *SCTransform* method (with variable.features.n=2000, conserve.memory=T, Seurat package version 4^45^), dimensionality reduction by PCA (Seurat *RunPCA*, npcs=30), construction of k-NN neighbor graph (Seurat *FindNeighbors*, dims=1:30) and Louvain community detection clustering (Seurat *FindClusters*, resolution=0.2, algorithm=1).

#### Automatic classification of cell types

We automatically classified nuclei into one of the following eight major cell types: excitatory neurons, inhibitory neurons, astrocytes, microglia, oligodendrocytes, OPCs, endothelial, and pericytes. The automatic annotation of nuclei was done by a weighted ElasticNet-regularized logistic regression classifier, fitted over our previous atlas of the human aging DLPFC from 24 individuals^4^ with a total of 182,739 nuclei (**Supplementary Fig. 1c**). The gene count matrix of the previous atlas^5^ was log normalized (Seurat *NormalizeData*) and scaled (Seurat *ScaleData*, method=vst) over the top 700 variable features (Seurat *FindVariableFeatures*, excluding non-coding non-annotated loci with a pattern of ^(AL|AC|LINC)\\d+).

To select the optimal regularization parameter we applied 10-fold cross-validation (*cv.glmnet* method, glmnet package^46, 47^) over randomly selected 75% of the data. To ensure the capture of rare cell types such as pericytes, we weighed samples as 1/nk for the number of nuclei of the cell-type present in the training set. We selected the ElasticNet mixing parameter of *α*=0.25 (to increase the sparsity of the fitted model) by evaluating test accuracy over the remaining 25% of the data. The fitted model used only 121 features and achieved a test accuracy of 99.95.

#### Removal of low quality cells

Low-quality nuclei were identified by the total number of UMIs (nUMI) and the number of unique genes (nGene). As different brain cell types have inherently different RNA quantities, we learned cell-type specific thresholds over these parameters. Thresholds were optimized based on hand annotation of 10 pooled libraries and applied to all 128 libraries to classify low-quality cells, and remove such cells from the downstream analysis (**Supplementary Fig. 1d**). Clusters of the 10 pooled libraries were manually curated to low- and high-quality clusters based on the nUMI and nGene distributions (Seurat *VlnPlot*). Then, we selected the cell-type specific thresholds as the median of all optimal nUMI and nGene parameter pairs, scored using the harmonic mean of the precision and recall. nUMI and nGene thresholds were: excitatory neurons 2232 and 1916; inhibitory neurons 800 and 100; astrocytes 800 and 616; microglia 400 and 253; oligodendrocytes 400 and 253; OPCs 695 and 253; vascular cells 400 and 253; and pericyte cells 400 and 100 respectively (**Supplementary Fig. 1e,f**). Low-quality clusters were removed as well, by a Soft-SVM classifier fitted over the 10 pooled libraries and using the (1) proportion of nuclei annotated as low quality (by nUMI and nGene threshold); (2) average entropy of cell-type prediction, and (3) proportion of doublets by the demuxlet algorithm.

#### Doublet detection

Between sample doublets were identified by the demuxlet algorithm, based on the sample barcodes (**Supplementary Fig. 1g**). Within sample doublets were predicted *in silico* based on their RNA profiles. To predict doublets, we ran DoubletFinder^48^ (*DoubletFinder_v3* method, pN=0.5, pK=75/(1.5*(#nuclei in library)), nExp=0, sct=T) over each of the libraries (**Supplementary Fig. 1h**). Thresholds for DoubletFinder predictions were determined separately for each library, based on the maximal Matthew’s Correlation Coefficient (MCC) compared to the demuxlet-identified doublets (**Supplementary Fig. 1i**). Further, as DoubletFinder is not designed to identify doublets of the same cell type, we modified it to simulate doublets of parent nuclei of different cell types, inferred based on the cell-type classification (https://github.com/GreenGilad/DoubletFinder). By high-resolution clustering of the nuclei (Seurat *FindClusters*, resolution=1.5) we expanded and marked as a doublet any nuclei predicted to be a demultiplexed doublet, a DoubletFinder doublet, or belonging to a cluster consisting of more than 70% DoubletFinder doublets.

#### Unified UMAP space

To compute a UMAP embedding consisting of all nuclei, we created a Seurat object over a subset of nuclei and projected the remaining nuclei onto the UMAP space computed for the subset. Since at the time of writing Seurat did not support 64-bit matrices, we were unable to create a single 1.63 million nuclei large object. We randomly sampled 30 out of the 128 libraries and created a single Seurat object consisting of 400,000 nuclei. We then followed a similar downstream analysis of normalizing, scaling and PCA- and UMAP embedding as described above (using 4,000 variable features and a PCA space of 50 dimensions). These PCA- and UMAP spaces are referred to as reference spaces.

The remaining nuclei were then projected as follows. Each library was normalized and scaled using only the variable genes used by the reference (Seurat, *SCTransform*, specifying residual.features argument as the reference variable genes). Then, the scaled data was projected onto the reference PC space using the reference’s feature loadings. Now that both sets are embedded in the same PC space, the remaining nuclei were projected onto the reference’s UMAP space (Seurat *ProjectUMAP*, setting reference- and query reductions as “pca”).

### Sub-clustering analysis

Based on the cell-type classification, we partitioned nuclei into subsets of the different cell classes, and we performed the sub-clustering analysis separately per cell type (endothelial cells and pericytes were analyzed together as part of the vascular niche). For each cell class, we performed the following analysis steps: (1) removed genes expressed in fewer than 15 nuclei as well as non-coding and non-annotated genes (pattern=^(AC\\d+{3}|AL\\d+{3}|AP\\d+{3}|LINC\\d+{3})), (2) removed cells with more than 10% mitochondrial RNA (Seurat *PercentageFeatureSet*, pattern=‵^MT-‵), and (3) removed any residual doublet cells that DoubletFinder did not detect. In the case of oligodendrocytes and OPCs, we did not remove nuclei predicted as oligodendrocyte-OPC doublets.

We iteratively ran the normalization and clustering pipeline as described for each library to remove additional low quality or doublet nuclei, with the following parameters: (1) *SCTransform* method with variable.features.n=4000 (for most cells) and n=2000 in the vascular niche subset, (2) *RunPCA* with npcs=50, (3) *FindNeighbors* with dims=1:50, *FindClusters* with resolution=1.5, algorithm=4 (Leiden), method=igraph, and (4) *RunUMAP* with dims=1:50, min.dist=0.1. After each such run, we removed clusters with low-quality cells or a high percentage of doublet cells (following the guidelines described above). To validate the doublet annotations we plotted the expression level of canonical cell-type markers, and small clusters expressing multiple markers of multiple cell types were also removed.

After cleaning the nuclei per cell type, we performed sub-clustering analysis per cell type. For each cell typethe sub-clustering was performed over multiple resolutions (Seurat *FindClusters*, algorithm=4, method=igraph), and the resolution was determined by the (1) differential gene expression per cluster (Seurat, *FindAllMarkers*, test.use=negbinom), (2) functional annotations of the differential signatures and (3) the proportions of clusters across individuals. Clusters that did not have differential genes or clusters specific to a single individual were united with the neighboring cluster with a similar RNA profile.

Due to the high sequencing depth and number of nuclei, excitatory neurons were split into two major sets (one consisting of layer 2-4 pyramidal neurons, CUX2+ expressing nuclei, and the other consisting of all others). Cleanup and sub-clustering analysis was performed for each subset separately. Neuronal clusters at the borders were then re-clustered to avoid misclassifications due to the split.

### Differential expression and functional annotation

Within a given cell class subset, DEGs were computed between each cluster and the rest using a negative binomial test and controlling for the post-mortem interval and participants’ batch (Seurat *FindAllMarkers*, test.use=negbinom, latent.vars = batch, pmi). In the case of excitatory neurons, as we were not able to create a single Seurat object to facilitate all cells, we partitioned genes into 6 groups. We then merged CUX2+ and CUX2-cells taking a single group of genes at a time and ran the negative binomial test as above. After merging the results, we corrected for multiple hypothesis testing in the same manner as is performed by the Seurat functions.

The functional annotation of clusters was inferred by (1) identifying up- and downregulated gene pathways and (2) grouping them based on their similarities. We identified up- and downregulated pathways using clusterProfiler^49^ calling *compareCluster* method (formula=id∼cluster+direction) using KEGG (fun=enrichKEGG, organism=hsa), Reactome (fun=enrichPathway) and GO (fun=enrichGo, OrgDb=org.Hs.eg.db, ont=BP,MF,CC) annotations, while setting the background universe to the set of all genes present in the data. Redundancy between Go terms were removed (*simplify* method, clusterProfiler).

To better understand the functionality captured by the pathways, we clustered the inferred upregulated or downregulated pathways of a given subpopulation. Given a list of pathways, we first computed two one-hot encoding matrices of pathways over genes: an “evidence” matrix and a “prior” matrix, both sharing the same rows (pathways). The columns of the evidence matrix are the union of all DEGs found for the particular subpopulation and take a value of 1 at place *i*,*j* if and only if DEG *j* is associated with pathway *i*. The columns of the matrix are the union of all genes associated with the pathways of the particular subpopulation, and take a value of 1 at place *i,j* if and only if gene *j* is associated with pathway *i*. Both matrices were then converted into pathways pairwise similarity matrices (*term_similarity* method, simplifyEnrichment package, method=kappa). We used the kappa coefficient as the measure of similarity to take into account the chance of a given gene to be associated with two different pathways. The partitioning of the pathways was done over the joint adjacency matrix being the element-wise average of the two similarity matrices, using the binary-cut algorithm (*cluster_terms* method, simplifyEnrichment package, method=binary_cut). The partitioning cutoff passed to the algorithm was adjusted manually based on the visual inspection of the joined adjacency matrix.

### Computation of subpopulation proportions

We divided cell subpopulations into 7 cell types: excitatory neurons, inhibitory neurons, astrocytes, microglia (including monocytes and macrophages), oligodendrocytes, OPCs and vascular niche composed of vascular cells, pericytes, SMC and fibroblasts. Rare subpopulations of erythrocytes, CD8+ T cells, NK cells and neutrophils, for which we have low total abundances, were not included. We then calculated for every participant the subpopulation proportion *within* the relevant cell type represented by matrix *P* whose rows represent participants and columns represent subpopulations:

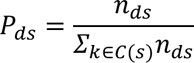

for *d* a participant, *s* a subpopulation *C*(*s*) the set of all subpopulations in the cell type of *s* and *n_ds_* the abundance of subpopulation *s* in participant *d*. The row [*P*]_*d*_ is the cellular environment of participant *d*, while column [*P*]_.,*s*_ is the vector of subpopulation proportions for subpopulation *s* across all participants. The rows and columns of *P* were used for association- and causality analyses as well as the BEYOND analysis (see sections below).

### Statistical analysis associating subpopulations to AD-related traits

Statistical associations between traits and subpopulations were tested by regressing traits on the square-root proportions of the subpopulation: 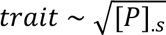 for subpopulation *s* and *P* the proportions matrix (defined in the section above). To remove potential confounding effects, we adjusted for (1) age at death, sex and post-mortem interval, and (2) snRNA-seq library quality reflected by the number of cells captured and total genes detected for each participant. Results were corrected for multiple hypothesis testing by calculating the false discovery rate (FDR, *p.adjust*, method=BH) within each tested trait. We applied a similar approach when associating CelMod predicted subpopulation proportion (see section below) and AD-related traits, adding RIN scores as the measure of library quality. To integrate the statistical associations calculated over the snRNA-seq and the bulk-predicted datasets we applied a meta-analysis approach using a weighted z-statistic^50^

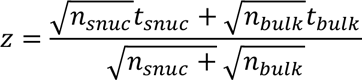

Meta-analysis results were then corrected for multiple hypothesis testing (*p.adjust*, method=BH).

#### Inferring cell-state proportions from bulk RNA-seq

We leveraged bulk RNA-seq data generated from DLPFC of 1,092 samples. As described in Lee *et al.*,^51^ RNA-seq data was generated in three sessions: 1) 10 batches of 739 subjects were sequenced at the Broad Institute, using Qiagen’s miRNeasy Mini Kit and RNase-free DNase set. RNA was quantified using Nanodrop and RNA quality evaluated using Agilent’s Bioanalyzer. Samples with RIN score >5 and quantity threshold >5 micrograms were submitted for library construction. Sequencing was conducted using the Illumina HiSeq2000 with 2×101bp reads for a targeted coverage of 50 million paired reads. 2) 2 batches of 124 subjects were sequenced at the New York Genome Center, using the KAPA Stranted RNA-seq kit with RiboErase. Sequencing was carried out on the Illumina NovaSeq6000 using 2×100bp reads targeting 30 million paired reads per sample. 3) One batch of 229 samples were sequenced at the Rush Alzheimer’s Disease Center, using the Chemagic RNA tissue kit, with RNA Quality Number calculated using the Fragment Analyzer. 500ng total RNA was used to generate the RNA-seq library, after rRNA depletion with RIbogold. Libraries were sequenced on the Illumina NovaSeq 6000 with 2×150bp reads at 40-50 million reads per sample.

RNA-seq reads were aligned to the GRCh38 human genome and gene-level counts were normalized to transcripts per million (TPM). Outlier samples were removed based on MDS plots and quantiles of expression profiles, and genes with median TPM<10 were filtered out to reduce technical noise. Log2-transformed TPM values in the expression matrix were included in a linear regression model with the following covariates: age at death, sex, batch, library size, percentage of coding bases, percentage of aligned reads, percentage of ribosomal bases, percentage of UTR bases, median 5’ to 3’ bias, median CV coverage, post-mortem interval, and study index (ROS or MAP).

We used the CelMod package^4^ (https://github.com/MenonLab/Celmod) to infer cell subpopulation proportions from the residuals of the bulk RNA-seq data, for participants without snRNA-seq profiles. As described previously, CelMod uses a consensus regression model trained on matched snRNA-seq proportions and bulk RNA-seq to extrapolate cell subpopulation proportions to samples with only bulk RNA-seq available. Briefly, the approach builds a regression model for each bulk RNA-seq gene using the cell type proportions as the predictor variable. Genes are then ranked by goodness-of-fit for each cell type, and the mean of the top genes is used as the prediction for the proportion of a given cell type in samples having only bulk data. The only free parameter – the number of genes used in the consensus – is determined by a 5-fold cross-validation of the training set.

With a total of 408 overlapping participants between the snRNA-seq and bulk RNA-seq samples, we fitted a CelMod model using 306 (75%) participants. The remaining 102 participants were kept as the hold-out set to assess the reliability of the predictions. Because of the distribution of snRNA-seq proportions, we applied CelMod to the square root of the proportion values. After running CelMod on each major cell class separately, we calculated the Spearman correlation of the predicted proportion versus the snRNA-seq proportion on the holdout set. This per-cell type Spearman correlation provides an estimate of the reliability of the prediction for each cluster. Only sub-populations for which correlation was positive and significant (FDR<0.05) were considered reliable and used in downstream analysis.

### Causal Modeling

In this section, we focused on the most robust subpopulation – AD trait associations by selecting those with the association FDR<0.01 with at least two tested AD traits: Mic.12, Mic.13, and Ast10. We first assessed the association of *APOE*ε4, the strongest genetic risk factor of AD, with the identified key cell states. Subsequently, we tested whether AD endophenotypes (starting with Aβ) mediate the observed *APOE*ε4 – cell state association. This step uses *APOEε4* as the genetic anchor that guides the direction of effect, leveraging the fact that the genetic variation is not subject to reverse causation^52^. Then, we built on the widely accepted causal chain of AD pathogenesis that starts with Aβ, which leads to neocortical tau accumulation and cognitive decline^26–28^, and performed a series of causal mediation analyses to determine the most plausible location of each cell state in the core sequence of AD (Aβ->tau->cognitive decline). In mediation models, we used AD endophenotypes and square-rooted cell state proportions as continuous variables and performed causal mediation analyses using a non-parametric bootstrap option with 10,000 simulations in the R *mediation* package^53^. Then, we constructed a structural equation model (SEM) based on the mediation analyses using the R *lavaan* package^54^. We first calculated the residuals of each variable by regressing age, sex, PMI, the estimated number of cells (per individual), and the number of detected genes (per individual). An exception was cognitive decline, as this variable has already been residualized against demographic covariates (random slope from linear mixed effect model, adjusting for baseline age, sex, and years of education)^39^. We did not allow cognitive decline to be the parent (causal) node, as cognitive decline is a symptom rather than the cause of AD pathophysiology and cellular environment changes. To replicate our findings in independent individuals, we repeated the same procedure, using the CelMod-predicted cell subpopulation proportions data from ROSMAP participants who did not contribute to the quality-controlled snRNA data (Replication sample).

### BEYOND analysis

To implement the BEYOND strategy, we used the cellular landscape matrix *P* ∈ [*0*,*1*]^424×92^ (defined above), where each row [*P*]_*d*_ represents the cellular environment of participant *d*. We leveraged our large sample of individuals to learn the *cellular landscape* manifold as captured by the columns of *P*. For convenience we stored *P* as an AnnData^55^ object.

#### Step 1: Visualizing the cellular landscape manifold

We embedded the cellular environments using three manifold learning algorithms: (1) PHATE^56^ (scanpy package, *external.tl.phate* method, k=10, n_components=2 and 3, a=40, knn_dist=cosine, mds_dist=cosine, mds_solver=smacof); UMAP^57^ (scanpy package *tl.umap* method, maxiter=3000, alpha=0.1, min_dist=0.25); and tSNE^58^ (scanpy package *tl.tsne* method, n_pcs=0, use_rep=”X”, learning_rate=100).

#### Step 2: Pseudotime analysis

Inference of pseudotime and trajectories was done using Palantir^30^. In short, Palantir projects the input data onto a multi-dimensional diffusion space and constructs a neighboring graph. It then iteratively refines the shortest paths from a user-defined starting point to each given data-point, defining this relative distance as the pseudotime. Lastly, it constructs a Markov chain using the neighboring graph and inferring directionality by the pseudotime. Palantir defines trajectories as the terminal states of the Markov chain and trajectory probabilities as the probability of a given data-point to reach each of the terminal states.

We ran Palantir over the cellular landscape excluding participants of clusters 9 and 10, and setting the starting point as the medoid point of clusters 1 and 3 (Palantir *run_diffusion_maps* and *determine_multiscale_ space* methods, n_components=5, knn=50; Palantir *run_palantir* method, knn=30, max_iterations=100, scale_components=F, n_jobs=1, use_early_cell_as_start=F).

#### Step 3: fitting and plotting dynamics

Dynamics were computed by regressing the feature values over the pseudotime in a specific trajectory using a generalized additive model (GAM), similarly to the strategy used by Palantir^30^. Features used were participants’ traits, subpopulation proportions or community proportions. We then used the fitted model to predict the final dynamics over equidistant pseudotime values. In detail, we spline-fitted a GAM for feature *y* in trajectory *j*:

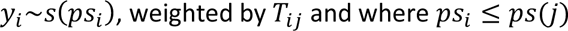

where *i* a participant, *j* a trajectory, *ps*_I_ the pseudotime of participant *i*, *ps*(*j*) the terminal pseudotime of trajectory *j* and *T*_Ij_ the trajectory probability matrix of participant *i* in trajectory *j* (mgcv *gam* method^59^, formula=*y*∼*s*(*ps*), weights=*T*_.j_). The final dynamics were predicted over equidistant pseudotime values in the range [0, *ps*(*j*)] (mgcv *predict.gam* method, se.fit=TRUE). When plotting the dynamics we presented the predicted feature values over the equidistant pseudotime values, as well as a confidence interval area of predicted value ±2×se, as retrieved from *predict.gam*.

#### Step 4: Construction of cellular communities

Partitioning of subpopulations into cellular communities was done using two sorts of measures: (1) similarity in subpopulation dynamics and (2) co-occurrences of subpopulations across participants. Similarities of dynamics were calculated using a weighted adaptive RBF kernel over the z-scored dynamics matrix, computed as follows: we represented each subpopulation *s* by its dynamics along both trajectories 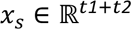 for *t1* and *t2* the number of equidistant pseudotime values used for the predicted dynamics (see section above). We obtained 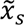 by centering and standardizing 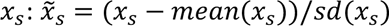. We then computed *M* the Mahalanobis distance between every two subpopulations as *M_sk_* = (*x_s_* − *x_k_*)^Τ^*W*(*x_s_* − *x_k_*) for *W* a diagonal matrix whose first *t1* diagonal values are *1*/*t1* and the rest *1*/*t2* (i.e. equally weighing both trajectories). Lastly, we calculated the adjacency matrix as *A_dyn_* = *exp*(−*M*/σσ^Τ^), where the division is performed element-wise and clipped to zero values smaller than 10^-4^. σ is the vector of local densities at each subpopulation calculated as 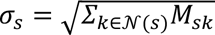 and 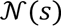 is the set of 5-nearest neighbors of *s* according to *M*. The co-occurrence matrix *A_co_* was calculated as the pairwise spearman correlation matrix of subpopulation proportions across all participants (*cor* method, stats package, use=pairwise.complete.obs, method=spearman). That is, [*A_co_*]_*sk*_ = *cor*([*P*]_.*s*_, [*P*]_.*k*_) for *P* the proportions matrix defined in the sections above.

We used the Leiden community detection algorithm over the multiplexed 3-layered graph induced by the matrices *A_dyn_*, *A_co_* after zeroing negative correlations and −*A_co_* after zeroing positive correlations (*optimise_partition_multiplex* method, leidenalg python package^60, 61^), with layers weighted as 1,1,-1 respectively. We used the RBConfigurationVertexPartition partitioning model^62, 63^, specifying the resolution parameter for each layer as the value maximizing the modularity (*resolution_profile* method, leidenalg python package, partition_type=RBConfigurationVertexPartition, resolution_range= [10^-2^, 10], number_iterations=-1). Based on visual inspection of the partitioning, community #0 was further partitioned into communities #2 and #3 using the same procedure as above over the subgraph induced by community #0.

Once subpopulations were partitioned into cellular communities, we assigned participants with a community proportion by averaging the normalized subpopulation proportions, and further normalization such that community proportions for every participant sums to one. Community dynamics along trajectories was calculated using the same procedure as used for traits- or subpopulation-dynamics.

### smFISH quantification and analysis

#### RNAscope experiment

RNAscope Multiplex Fluorescent Reagent Kit v2 (ACD, 323100) was used to perform the RNAscope experiments on 13 individuals from the New York Brain Bank (4 cognitively unimpaired, no AD; 2 mild cognitive impairment, MCI; 7 AD). 6 μM paraffin-embedded tissue sections were deparaffinized with CitriSolv Clearing Agent (Decon Laboratories, In., 1601) for 20 min at Room Temperature (RT), followed by an ethanol series (100%, 100%, 70%) for 30 seconds each. The slides were then put in distilled water for 1 minute at RT. Then, slides were incubated with hydrogen peroxide for 10 min at RT, then washed with distilled water twice to stop the hydrogen peroxide reaction. Antigen retrieval was performed with pH 6.0 citrate (Sigma-Aldrich, C9999) and heating with a microwave for 25 min at 400 watts. After 5 minutes in tap water, slides were immersed in 100% ethanol for 1 minute. Once the slides had fully dried at RT, the Super Pap Pen Liquid Blocker (Newcomer Supply, 6505) was used for drawing a hydrophobic barrier around the tissue section. Slides were blocked for 30 minutes at RT with RNAscope Co-Detection Antibody Diluent.

The slides were stained with IBA1, a microglia marker (Wako, 01127991) diluted in RNAscope Co-Detection Antibody Diluent at 1:50 and incubated for 2 hours at RT. Slides were then washed with PBS-T (PBS + 0.1% Tween-20) and then submerged in 10% neutral buffered formalin (Sigma-Aldrich, HT5011) for 1 hour at RT. After washing with PBS-T, slides were treated with RNAscope protease plus and incubated for 40 min at 40 °C in the RNAscope HybEz II oven. After washing with distilled water, RNAscope probe mix was added to the slides and incubated for 2 hours at 40 °C. Finally, the slides were washed with the RNAscope wash buffer and incubated with 5X SSC (Sigma-Aldrich, S6639-1L) overnight at RT. The next day, the slides were washed with RNAscope wash buffer and incubated with AMP1 for 30 min at 40° C. Slides were washed twice with wash buffer and AMP2 was added and incubated for 30 min at 40 ° C. After washing, AMP3 and incubated for 15 min at 40° C. After 15 min of incubation at 40 °C with HRP-C1, slides were washed and Opal 570, diluted in TSA diluent buffer 1:700 was added and incubated for 30 minutes at 40° C. Slides were then washed and finally, an HRP blocker was added for 15 min. This HRP/Opal/block process was repeated using HRP-C2, HRP-C3, and HRP-C4 depending on the channel of the original probes, and Opal 690) dye. TSA-DIG reagent was used after HRP-C4 and incubated for 30 min at RT then blocked with HRP blocker and washed with PBS-T. The secondary antibody (ThermoFisher Scientific, A11055) for the IBA1 staining was diluted in Co-Detection Antibody Diluent at 1:500 and then incubated for 30 min at RT. After washing with PBS-T, slides were incubated with Opal Polaris 780 dye prepared in the Antibody Diluent/Block at 1:700 for 30 minutes at RT. Slides were washed with PBS-T and the lipofuscin autofluorescence was quenched using Trueblack (Biotium, 23007) for 2 min at RT and DAPI was added to stain the nucleus. All slides were mounted using the Prolong Gold (Thermo Fisher Scientific, P36934). The RNAscope probes used in this experiment were: TPRG1-C1 (ACD, 1047171-C1), MRC1-C3 (ACD, 583921-C3), and CPM-C2 (444811-C2).

#### Image signal quantification

For all slides, images were acquired using the Nikon Eclipse Ni-E immunofluorescent microscope at magnification X40, and approximately 30 pictures were acquired per subject. The captured images were analyzed using CellProfiler software. An extensive pipeline has been developed to automatically segment the microglia and detect transcripts (CPM, MRC1 and TPRG1) expressed by IBA1+ cells^12^. DAPI was defined as the primary object using the *IdentifyPrimaryObjects* module. For IBA1+ cells segmentation, the module *EnhanceOrSuppressFeatures* was used to enhance the IBA1 signal and to detect the ramification. IBA1+ cells were also defined as the primary object and using the *RelateObjects* module, only IBA1+DAPI+ cells were selected. To segment the transcripts signals (dots), we used the *EnhanceOrSuppressFeatures* module with “Speckles” as the feature type, and the *RelateObjects* module was used to relate the transcripts signals (CPM, TPRG1 and MRC1) to IBA1+ positive for DAPI cells. The morphology (eccentricity and compactness) of IBA1+DAPI+ cells was measured using the *MeasureObjectSizeShape* module and the intensity of each transcript as well as IBA1 was measured using the *MeasureObjectIntensity* module. The data was exported in excel format using *ExportToSpreadsh*.

#### Prediction of microglial subpopulation in smFISH data

We fitted a weighted logistic regression classifier to assign RNAscope observed cells with a microglial state (*cv.glmnet* method, glmnet package^46, 47^). The classifier was fitted over a randomly selected 80% of snRNA-seq microglia nuclei, with a design matrix of CPM, TPRG1 and MRC1 gene expression values, scaled to the ranges observed in the RNAscope data, and a response vector of microglial states with classes of Mic.12, Mic.13, Macrophages and Other for the remaining microglial states defined in the atlas. As the dataset is imbalanced towards the “Other” class, we weighed samples as 1/*n_k_* for *n_k_* the number of nuclei of state *k* present in the training set. We then used the fitted model to predict state probabilities for each of the RNAscope observed cells. Regarding state probabilities as a soft-assignment of states, we averaged the probabilities per participant to obtain subpopulation proportions, similar to the representation of the snRNA-seq data.

## Data and Software availability

All data and software will be available before publication.

## SUPPLEMENTARY TABLES

- Supplementary Table 1 - Clinicopathological characteristics of participants in discovery (snRNA-seq 465 participants) and replication (non-overlapping bulk RNA-seq, 864 participants) cohorts.
- Supplementary Table 2 - Atlas characterization - differential expression and pathway analysis of subpopulation vs. all other subpopulations, and pairwise analysis of subpopulation vs. subpopulation, for every cell type.
- Supplementary Table 3 - Endophenotype associations - subpopulation proportions, CelMod correlations and predicted proportions, snRNA-seq-, CelMod predicted- and meta-analysis for subpopulation-endophenotype associations.
- Supplementary Table 4 - smFISH results - smFISH marker expression, subpopulation predicted probabilities, endophenotype and morphological quantification, as well as subpopulation proportions per participant.

## Author contributions

P.L.D. and N.H. designed the study; C.M. prepared the single nucleus libraries and performed sequencing; M.F. performed the sequence alignment and demultiplexing analysis; G.G. performed the computational- and statistical analysis, with guidance of H.Y., M.F., N.H., V.M., P.L.D. and help from A.C., A.K. and C.W.; H.Y performed the causality modeling analysis; M.T performed smFISH experiment and image analysis; G.G., H.Y., N.H. and P.L.D. wrote the manuscript, and all co-authors edited it for critical comments. D.A.B. is PI of the parent ROS and MAP studies and obtained funding and performed study supervision. P.L.D. and D.A.B obtained funding for the project.

## Acknowledgements

We thank the individuals who have generously donated their brain to research through the Rush University Alzheimer’s Disease Center. The work was supported by: NIH RF1 AG057473 (P.D. & D.B.), U01 AG061356 (P.D. & D.B.), U01 AG046152 (P.D. & D.B.), R01 AG070438 (P.D.), R01 AG015819 (D.B.), U01 AG072572 (P.D. & P.S.), R01AG066831 (to V.M.); NIH K23 AG062750 (to H.Y.); CS-02018-191971; the Israel Science Foundation (ISF) research grant no. 1709/19, the European Research Council grant 853409, the MOST-IL-China research grant no. 3-15687, and the Myers Foundation (given to N.H.);

---

**Supplementary Figure 1:**
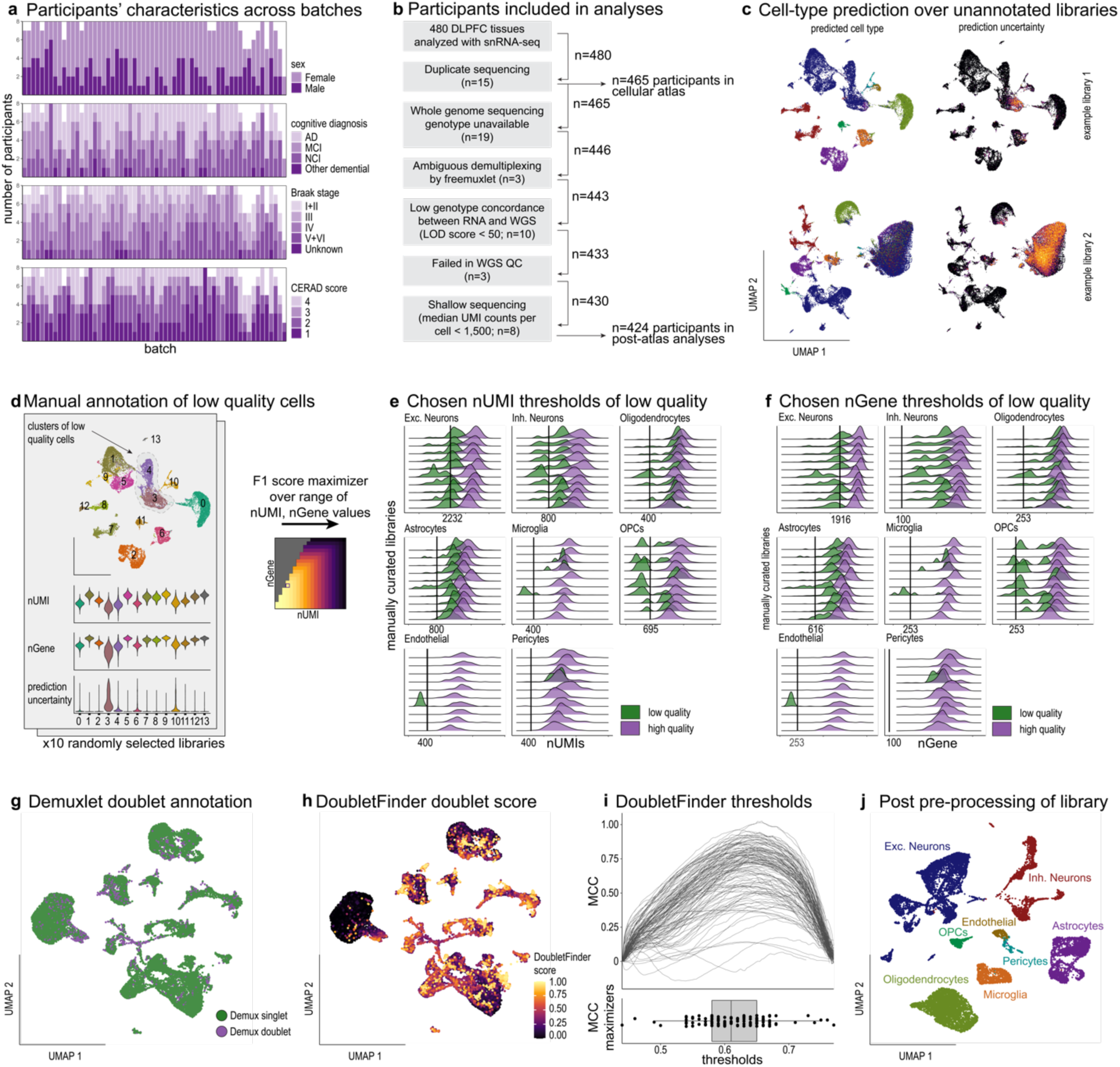
snRNA-seq libraries preprocessing and quality controls. (**a**) Batch design showing the distribution of sex, cognitive diagnosis, Braak stage and CERAD score for participants across the 60 experimental batches. (**b**) Overview of the filtering of snRNA-seq libraries. Starting with snRNAseq profiling for 480 participants, filtering of participants based on quality control measures retained nuclei from 465 participants used for building the cell atlas, and 424 participants with whole genome sequencing and sufficient number of nuclei for downstream analysis. (**c**) Automated cell-type classification. UMAP embeddings of example snRNA-seq libraries prior to the quality control steps, colored by the predicted cell type (left) or by the prediction uncertainty (Shannon entropy, right). (**d**) Manual annotation of low-quality clusters applied to 10 representative libraries to learn the low-quality threshold. Example of snRNA-seq library (top in c) manually curated for clusters of low-quality cells. Top: UMAP embedding, colored by cell type. Bottom: violin plots of the distribution per cell type of the number of UMIs (nUMI), the number of unique genes (nGene), and the cell type prediction uncertainty. (**e, f**) Cell type specific threshold for low quality cells. Distributions of (**e**) nUMI and (**f**) nGene in the 10 manually curated libraries. Black line: selected cell-type specific threshold. (**g**) Doublet detection by the demuxlet algorithm. UMAP embedding of example library (top in c) colored by doublet annotation obtained by demultiplex algorithm, which can detect doublets of nuclei from two different individuals based on their genetic diversity. (**Methods**). (**h**) Doublet detection by the DoubletFinder algorithm. UMAP embedding of example library (top in c) colored by the doublet-likelihood scores obtained by DoubletFinder, which can detect doublets of nuclei from two different cell populations based on their different RNA profiles. (**i**) Library specific threshold for doublet cells. Top: For each library, the Matthew’s Correlation Coefficient (MCC) of the DoubletFinder score compared to the demuxlet-identified doublets is plotted per DoubletFinder threshold (x-axis). Bottom: The library-specific threshold that maximizes the MCC. Dots: libraries of all experimental batches. Box: first and third quartiles; line: median; whiskers: outliers. (**j**) Post-processing library. UMAP embedding of example library (top in c), colored by the predicted cell type, after all preprocessing and quality control steps.

**Supplementary Figure 2:**
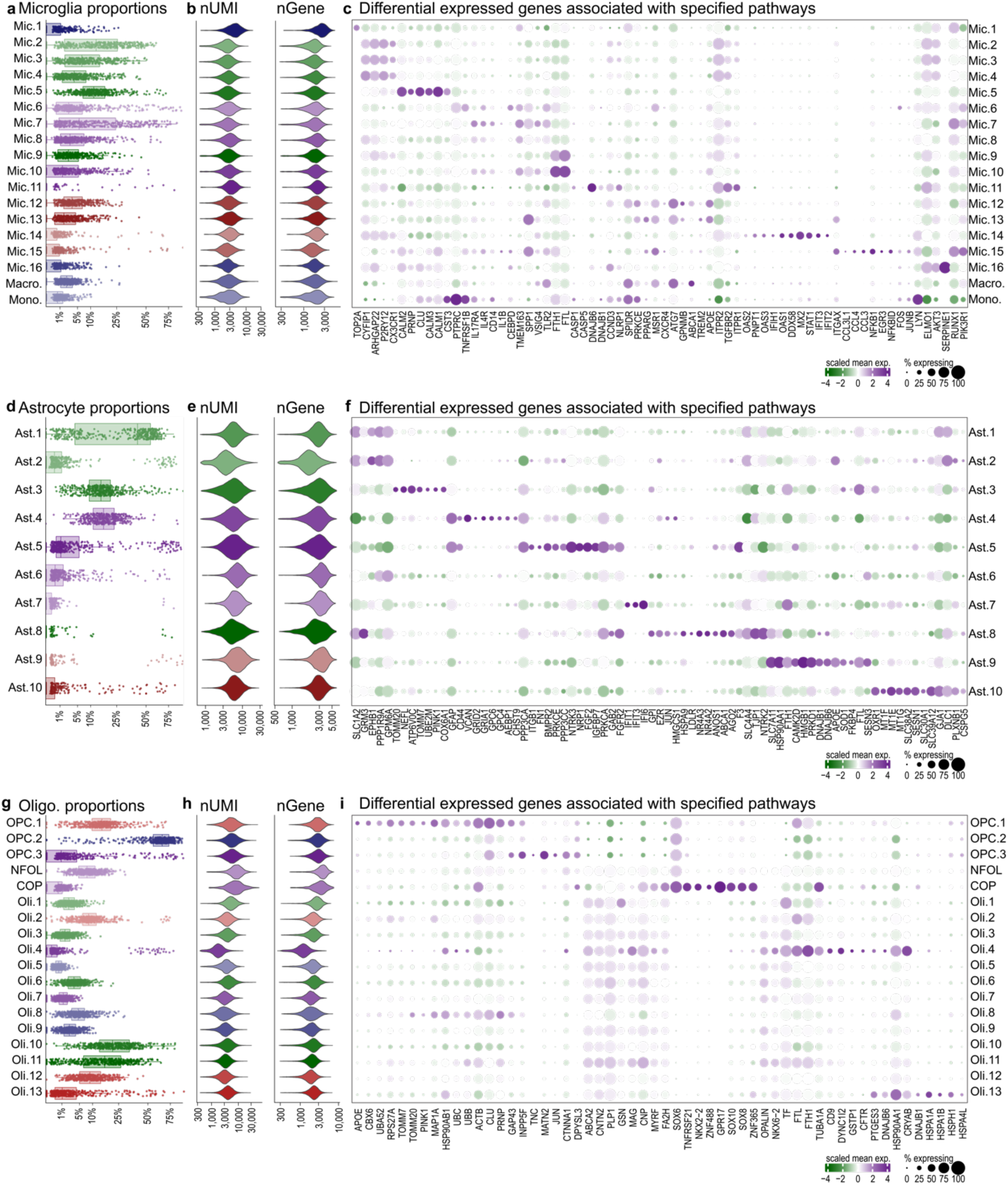
Quality controls and differential genes among glial subpopulations. (**a**,**d**,**g**) Subpopulation proportion across participants for (a) microglia, (d) astrocytes and (g) oligodendroglia cells. Dots: the subpopulation proportion scaled per individual. Box: first and third quartiles; line: median. (**b**,**e**,**h**) Distributions of number of UMIs (nUMI, left) and the number of unique genes (nGene, right) across subpopulations of (b) microglia, (e) astrocytes and (h) oligodendroglia. (**c**,**f**,**i**) Scaled gene expression (columns) of selected differential genes across cell subpopulations (rows) of (c) microglia, (f) astrocytes and (i) oligodendroglia. Dot color is the column scaled expression level of the expressing cells in the subpopulation, and size is the percentage of cells in the subpopulation expressing the gene.

**Supplementary Figure 3:**
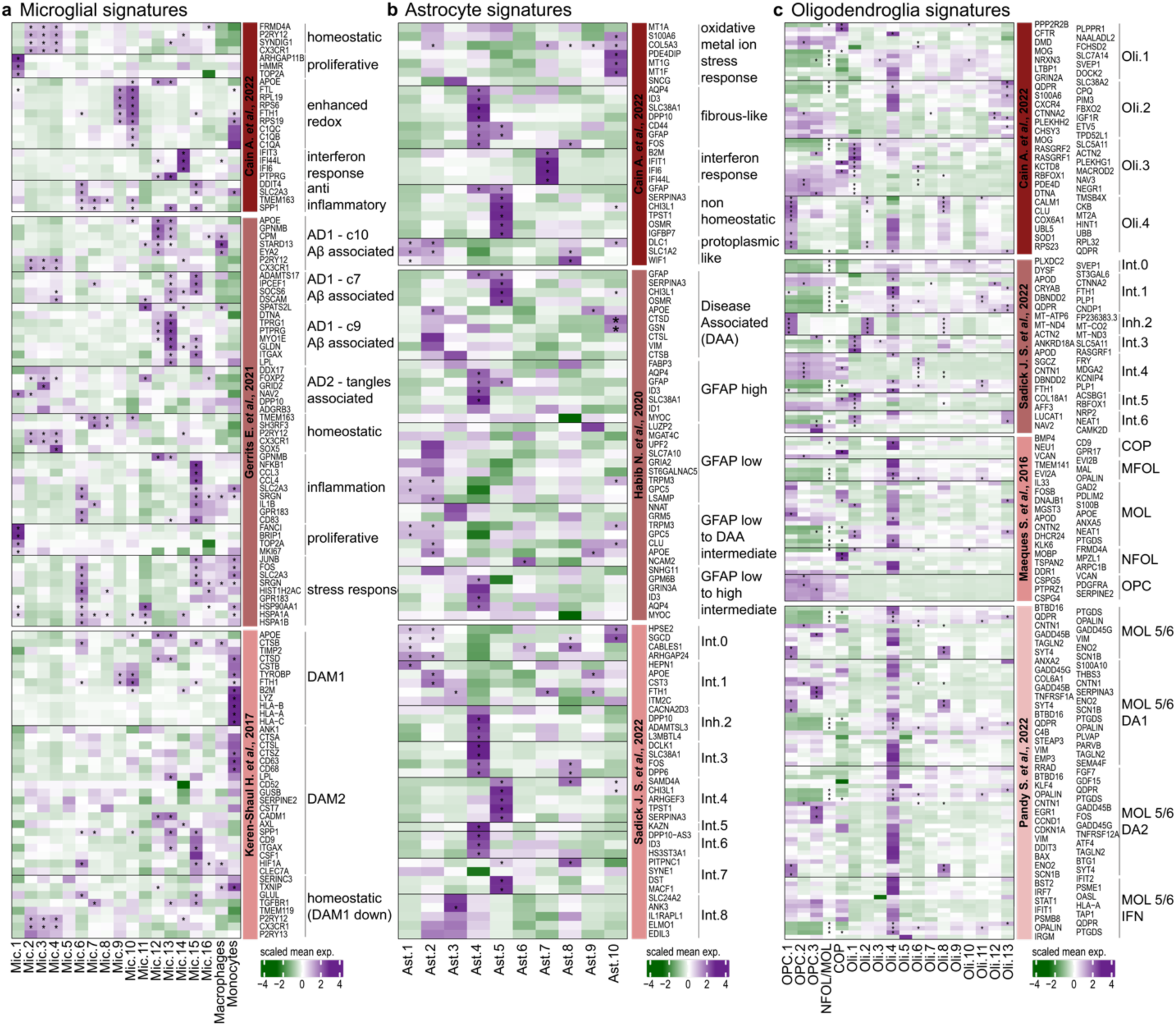
Comparison of glial diversity to the literature. Expression level of marker genes (rows) across glial subpopulations (columns) from published signatures of (**a**) microglia, (**b**) astrocytes and (**c**) oligodendroglia. Genes are separated by the published signatures and split by reference source. Color-scale: row scaled mean expression of expressing cells in the subpopulation. (*) marking significantly differentially expressed in the given subpopulation. Of note, genes defined in multiple signatures appear more than once.

**Supplementary Figure 4:**
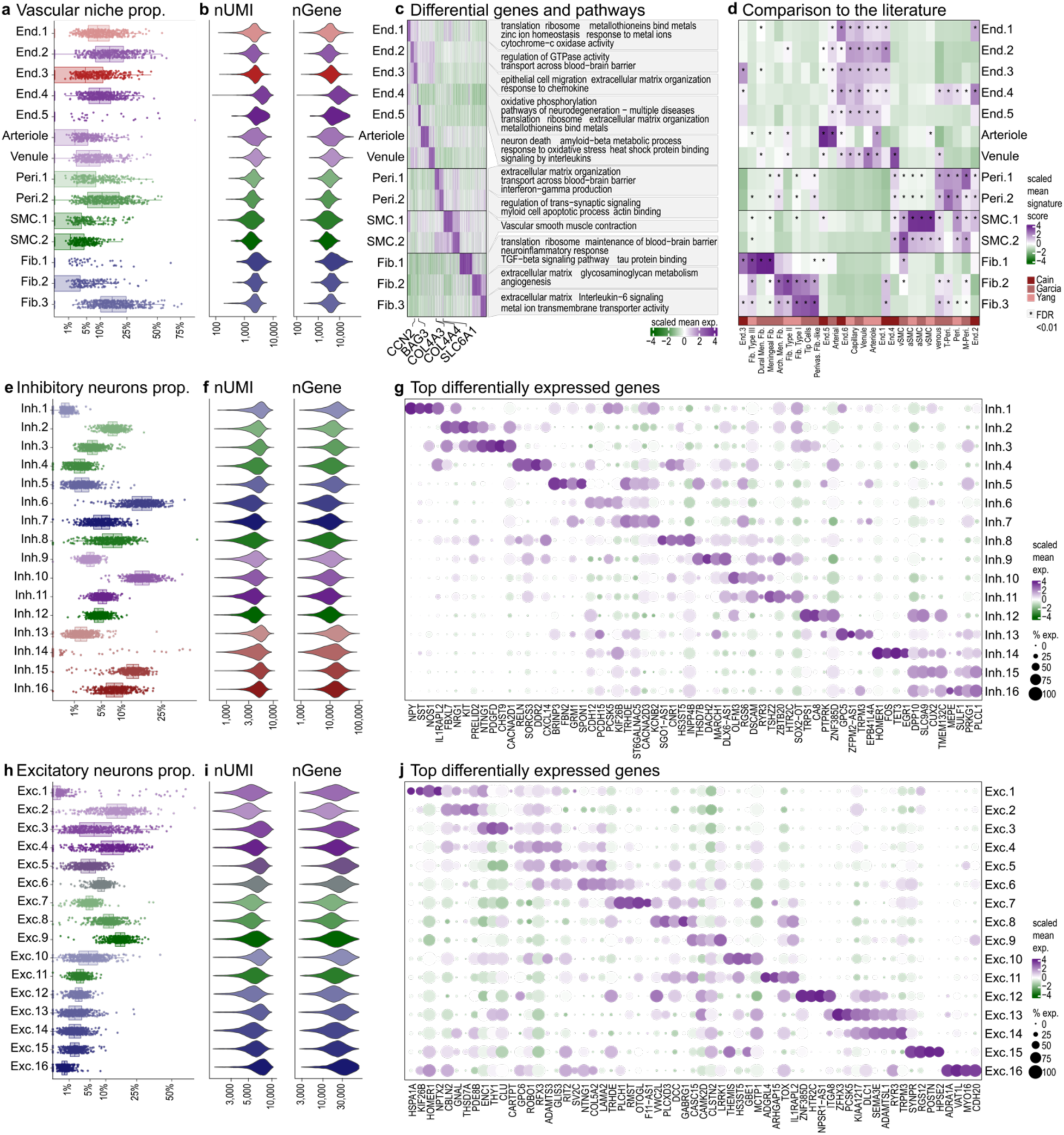
Vascular niche- and neuronal subpopulation diversity. (**a**,**e**,**h**) Subpopulation proportion across participants for (a) vascular niche, (d) inhibitory neurons and (g) excitatory neurons. Dots: the subpopulation proportion per individual. Box: first and third quartiles; line: median. (**b**,**f**,**i**) Distributions of number of UMIs (nUMI, left) and the number of unique genes (nGene, right) across subpopulations of (b) vascular niche, (f) inhibitory neurons and (i) excitatory neurons. (**c**) Distinct gene signatures and associated pathways across vascular niche cell subpopulations. Left: differentially expressed genes (columns) for different subpopulations (rows). Color scale: Scaled mean gene expression per cluster. Right: representative enriched unique pathways per subpopulation. (**d**) Comparison of vascular niche cell subpopulations to previous published signatures. Scaled mean signature score of published gene signatures (columns) within each subpopulation (rows)^4, 7, 15^. (*) = Significantly enriched signatures (U-test, FDR<0.01). (**g,j**) Differential gene signatures across neuronal subpopulations. Scaled expression levels for top differentially expressed genes (columns) for different subpopulations (rows) of inhibitory neurons (**g**) and excitatory neurons (**j**). Dot color is the column scaled expression level of the expressing cells in the subpopulation, and size is the percentage of cells in the subpopulation expressing the gene.

**Supplementary Figure 5:**
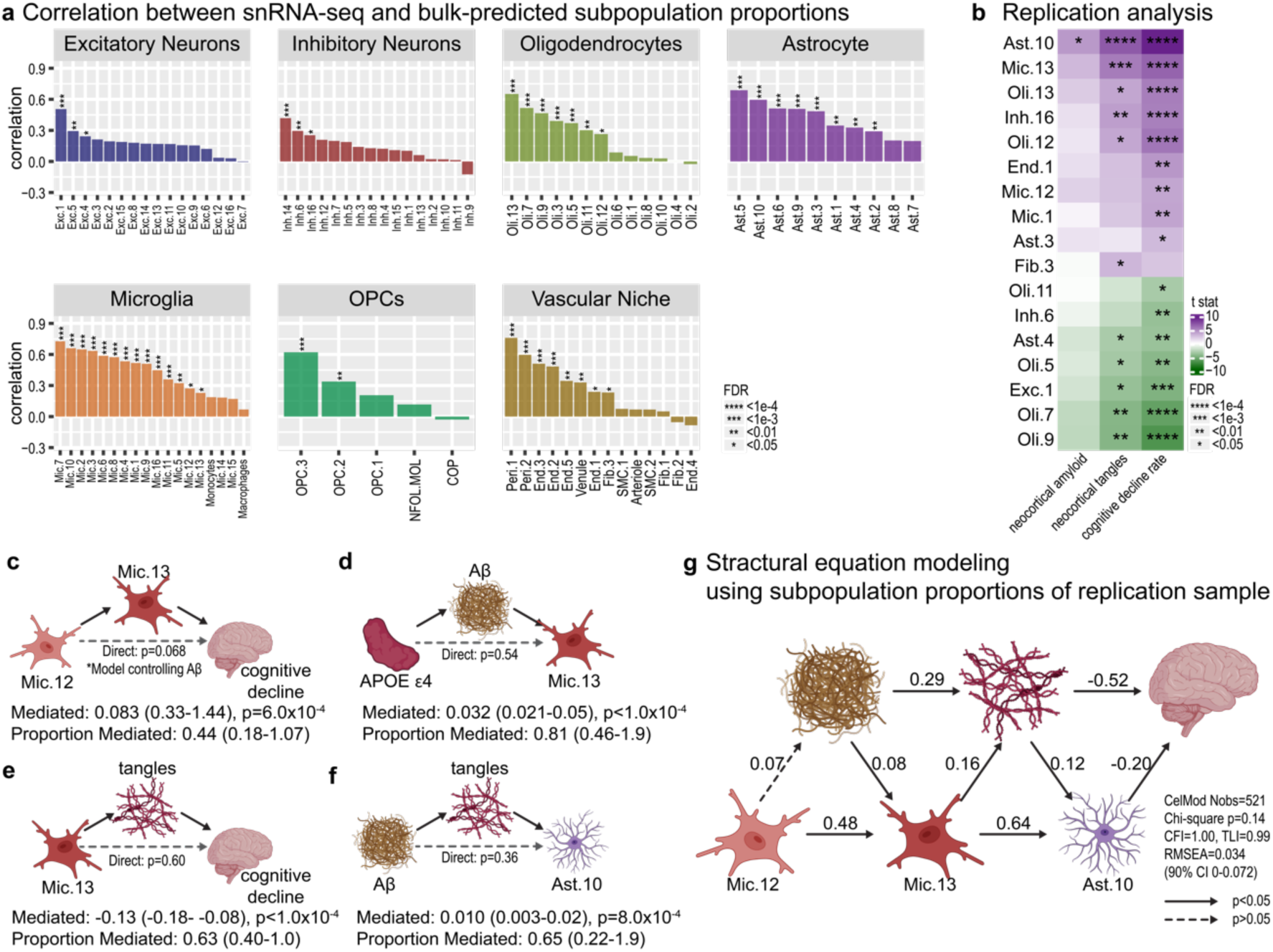
CelMod predictions of cellular compositions and mediation modeling. (**a**) CelMod predictions match snRNA-seq measurements. Spearman correlation between snRNA-seq proportions and CelMod bulk-predicted subpopulation proportions per cell type, over the test set of 102 participants (**Methods**). (**b**) *Replication analysis* associating subpopulations to AD traits: CelMod bulk-predicted subpopulation proportions significantly associated with at least one of the tested AD-related traits (linear regression with covariates FDR<0.05). n=684 participants. Color scale: association effect size indicating direction and strength from negative (green) to positive (purple) associations. (**c-f**) Causal mediation models positioning Mic.12, Mic.13 and Ast.10 within the AD cascade (accompanying results in **Fig. 3 f-i**). (**c**) Mic.13 mediates most of Mic.12 - cognitive decline association. (**d**) Aβ fully mediates APOE ε4 – Mic.13 association. (**e**) Tau mediates most of the Mic.13 – cognitive decline association. (**f**) Tau mediates most of the Aβ – Ast.10 association. (**g**) Replication analysis of structural equation model (SEM) positioning Mic.12, Mic.13 and Ast.10 within the AD cascade. We applied the final structural equation model (SEM; as done in Fig. 3j in the Discovery analysis) to the CelMod-predicted subpopulation proportions of participant of the Replication sample, with values for all tested AD traits (n=521). Numeric values indicate the relative strength of association. Solid arrows: Significant associations (p<0.05), Dashed arrows: Associations with p>0.05

**Supplementary Figure 6:**
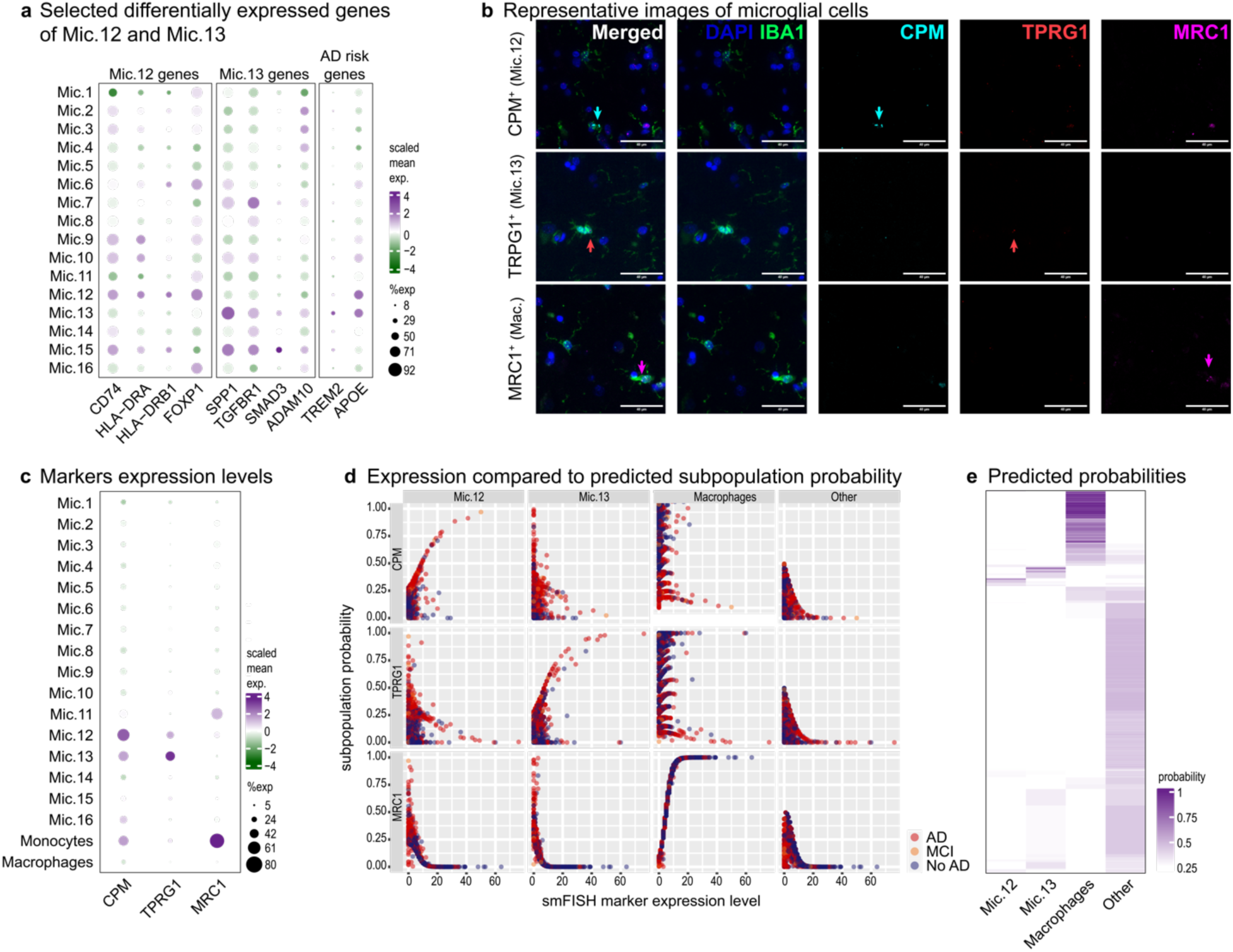
smFISH quantification analysis and validations for Mic.12 and Mic.13. (**a**) Marker gene expression across microglial subpopulations. Dotplot of gene expression (columns) across subpopulations (rows) for selected genes. Dots color is the column scaled mean gene expression level of all cells expressing the gene in the subpopulation, and dot size is the percent of cells in the subpopulation expressing the gene. (**b**) Representative RNA scope images. Split channels of representative RNA scope images shown in Fig. 4d, showing an example of a *CPM*^high^ (Mic.12 marker) microglia, a *TPRG1*^high^ (Mic.13 marker) microglia, and a MRC1^high^ (macrophage marker) cell. Microglia/myeloid cells (green, anti-IBA1 immunofluorescence), nuclei (DAPI), RNA scope probes targeting *CPM* (cyan, Mic.12 marker), *TPRG1* (red, Mic.13 marker), and *MRC1* (magenta, macrophage marker). (**c**) Expression of marker genes across subpopulations. Relative expression level of genes used for smFISH (columns) across microglia/myeloid subpopulations (rows). Dot plot as in a. (**d**) Subpopulation probability matches level of marker gene. Scatter-plot grid showing for each smFISH DAPI^+^IBA1^+^ cell its inferred probability (y-axis) for different microglia/myloid subpopulations (grid columns), over the expression level (x-axis) detected for each of the marker genes (grid rows). Showing the strong connection between each marker and its subpopulation probability: *CPM for* Mic.12, *TPRG1 for* Mic.13, MRC1 for macrophage. Dots are color coded by the clinical diagnosis of each individual. (**e**) Predicted probability of smFISH DAPI^+^IBA1^+^ as Mic.12, Mic.13, Macrophages or all other microglial cells.

**Supplementary Figure 7:**
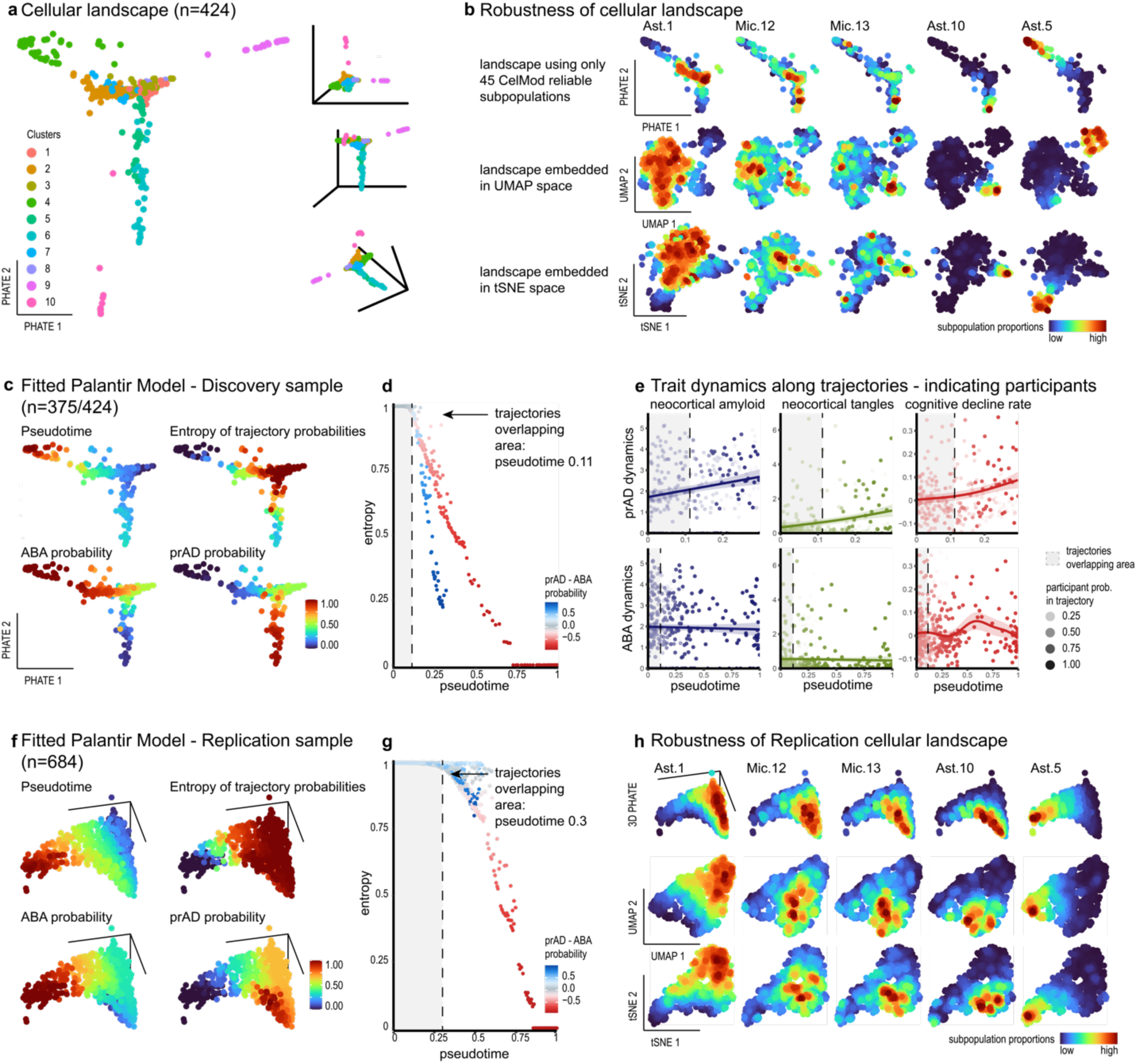
Robustness of the cellular landscape manifold of the aged neocortex. (**a**) The structure of the cellular landscape manifold of the aged neocortex. The cellular manifold captured by BEYOND algorithm for the discovery sample (n=424). Showing 2D and 3D PHATE^56^ embedding of each participant (individual dots) based on similarity of their cellular environments, represented as their cellular composition profiles in our cell atlas of 92 subpopulations (**Methods**). Outlier clusters 9 and 10 were excluded from the trajectories and dynamics analysis. (**b**) The cellular landscape manifold is robust to the embedding algorithm and number of subpopulations. Color scale: Locally-smoothen spatial distributions of subpopulation-proportions. Top: PHATE embedding using a subset of the subpopulations to represent cellular landscape; Center: UMAP embedding; Bottom: tSNE embedding. (**c-d**) Two cellular trajectories discovered in the aged neocortex. (c) Visualization of fitted Palantir trajectories model on the manifold (as in Fig. 5a) colored by the (clockwise from top-left): pseudotime, Shannon entropy of trajectory probabilities, ABA trajectory probability, and prAD trajectory probability. Values are not locally-smoothed. (**d**) Participant’s trajectory probabilities entropy drop along pseudotime. Each dot is an individual colored by the difference between the prAD and the ABA trajectory probabilities. The grey area indicates a pseudotime range (0, 0.11) in which the two trajectories are not well separated. (**e**) prAD and ABA trajectories have distinct trait dynamics. Trait-dynamics of AD-related traits along the pseudotime in each of the inferred trajectories (as in Fig. 5e), showing the datapoints (all individuals) used to fit the curves, shaded according to the trajectory probability. (**f-h**) Cellular landscape manifold and trait associations are validated in the replication sample. As c, d and b (respectively), but over the independent Replication sample of 684 participants with CelMod bulk-predicted subpopulations proportions. Using only the high confidence CelMod predictions for 45 subpopulations to represent cellular environments per participant (**Methods**).

**Supplementary Figure 8:**
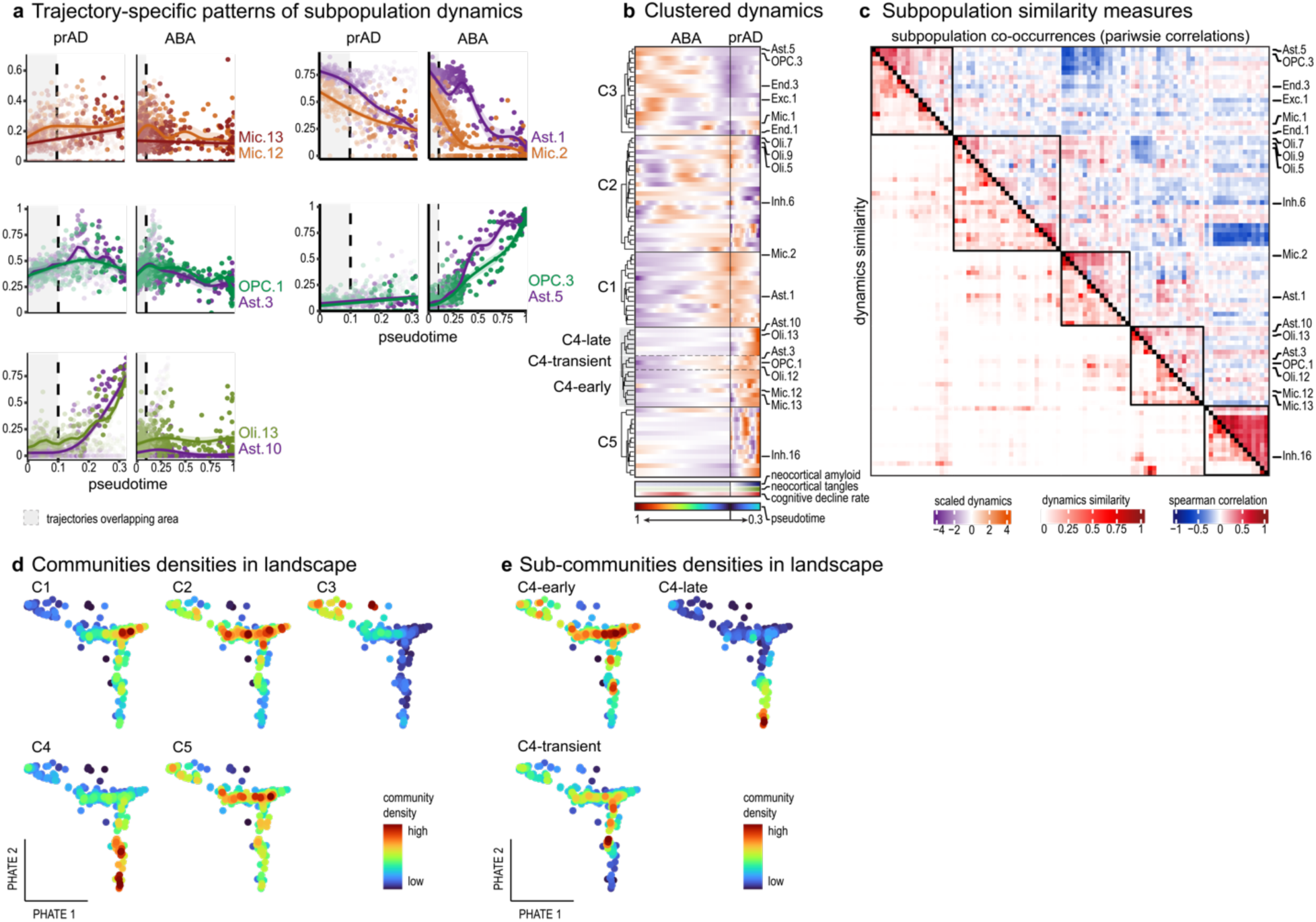
Cellular dynamics and multicellular communities in the aged neocortex. (**a**) Distinct dynamic patterns of cell subpopulations along the prAD and ABA trajectories. Patterns of subpopulation dynamics as shown in Fig. 6b including datapoints (all individuals) used to fit the curves, shaded according to the trajectory probability. (**b**) Clustering of subpopulations into multicellular communities. Scaled dynamics of each subpopulation along both trajectories. Clustering by BEYOND to communities. Bottom: dynamics of AD-traits. (**c**) BEYOND clustering of communities based on two measured of subpopulation similarities. Pairwise measures of subpopulation similarities used for clustering subpopulations into communities: showing subpopulations’ pairwise correlations (top) and dynamics similarity (bottom). Squares along the diagonal indicate the different cellular communities. (**d, e**) Distinct patterns of cellular community proportions along the cellular landscape manifold. Participants are colored by the locally-smoothed density of (d) communities and (e) sub-communities proportions embedded in the cellular landscape manifold (as in Fig. 5b).

